# Formicine ants swallow their highly acidic poison for gut microbial selection and control

**DOI:** 10.1101/2020.02.13.947432

**Authors:** Simon Tragust, Claudia Herrmann, Jane Häfner, Ronja Braasch, Christina Tilgen, Maria Hoock, Margarita Artemis Milidakis, Roy Gross, Heike Feldhaar

**Author notes:** Present address: General Zoology, Hoher Weg 8, Martin-Luther University, 06120 Halle (Saale), Germany (Simon Tragust).

## Abstract

Animals continuously encounter microorganisms that are essential for health or cause disease. They are thus challenged to control harmful microbes while allowing acquisition of beneficial microbes. This challenge is likely especially important for social insects with respect to microbes in food, as they often store food and exchange food among colony members. Here we show that formicine ants actively swallow their antimicrobial, highly acidic poison gland secretion. The ensuing acidic environment in the stomach, the crop, limits establishment of pathogenic and opportunistic microbes ingested with food and improves survival of ants when faced with pathogen contaminated food. At the same time, crop acidity selectively allows acquisition and colonization by Acetobacteraceae, known bacterial gut associates of formicine ants. This suggests that swallowing of the poison gland secretion acts as a microbial filter in formicine ants and indicates a potentially widespread but so far underappreciated dual role of antimicrobials in host-microbe interactions.

## Introduction

Animals commonly harbor gut associated microbial communities (Engel and Moran, 2013, Moran et al., 2019). Patterns of recurring gut microbial communities have been described for many animal groups (Brune and Dietrich, 2015, Kwong et al., 2017, Ochman et al., 2010). The processes generating these patterns are however often not well understood. They might result from host filtering (Mazel et al., 2018), a shared evolutionary history between gut associated microbes and their hosts (Moeller et al., 2016) involving microbial adaptations to the host environment (McFall-Ngai et al., 2013), simply be a byproduct of similar host dietary preferences (Anderson et al., 2012, Hammer et al., 2017), or result from interactions between microbes in the gut associated microbial community (Brinker et al., 2019a, García-Bayona and Comstock, 2018).

Food is an important environmental source of microbial gut associates (Blum et al., 2013, Broderick and Lemaitre, 2012, David et al., 2014, Hammer et al., 2017, Perez-Cobas et al., 2015) but also poses a challenge, the need to discriminate between harmful and beneficial microbes, as food may contain microbes that produce toxic chemicals or that are pathogenic (Burkepile et al., 2006, Demain and Fang, 2000, Janzen, 1977, Trienens et al., 2010). In social animals, control of harmful microbes in food while allowing the acquisition and transmission of beneficial microbes from and with food, is likely especially important. Eusocial Hymenoptera not only transport and store food in their stomach, the crop, but also distribute food to members of their colony via trophallaxis, i.e. the regurgitation of crop content from donor individuals to receiver individuals through mouth-to-mouth feeding (Gernat et al., 2018, Greenwald et al., 2018, LeBoeuf et al., 2016). While trophallaxis can facilitate the transmission of beneficial microbes, from an epidemiological perspective it can also entail significant costs, as it might open the door to unwanted microbial opportunists and pathogens that can take advantage of these transmission routes (Onchuru et al., 2018, Salem et al., 2015).

Here we investigate how formicine ants, specifically the Florida carpenter ant *Camponotus floridanus*, solve the challenge to control harmful microbes in their food while allowing acquisition and transmission of beneficial microbes from and with their food. Apart from specialized intracellular endosymbionts associated with the midgut in the ant tribe Camponotini (Degnan et al., 2004, Feldhaar et al., 2007, Russell et al., 2017, Williams and Wernegreen, 2015), formicine ant species have only low abundances of microbial associates in their gut lumen but carry members of the bacterial family Acetobacteraceae as a recurring part of their gut microbiota (Brown and Wernegreen, 2016, Chua et al., 2018, He et al., 2011, Ivens et al., 2018, Russell et al., 2017). Some formicine gut associated Acetobacteraceae show signs of genomic and metabolic adaptations to their host environment indicating coevolution (Brown and Wernegreen, 2019, Chua et al., 2020). But the recurrent presence of Acetobacteraceae in the gut of formicine ants potentially also reflects direct transmission of bacteria among individuals, selective uptake on the part of the ants, specific adaptation for colonizing ant guts on the part of the bacteria, or some combination of all three (Engel and Moran, 2013).

Generally, the immune system together with physiochemical properties of the gut environment maintains homeostasis between gut associated microbes and the host (Chu and Mazmanian, 2013, McFall-Ngai et al., 2013, Rakoff-Nahoum et al., 2004, Slack et al., 2009, Watnick and Jugder, 2020, Xiao et al., 2019, see also Foster et al., 2017). Highly acidic stomach lumens are ubiquitous in higher vertebrates, including amphibians, reptiles, birds and mammals (Beasley et al., 2015, Koelz, 1992), while in insects acidic regions have rarely been described from midgut regions (Chapman, 2013, Holtof et al., 2019). However in both, higher vertebrates and the fruit fly *Drosophila melanogaster*, acidic gut compartments together with the immune system serve microbial control and prevent infection by pathogens (Giannella et al., 1972, Howden and Hunt, 1987, Martinsen et al., 2005, Overend et al., 2016, Rakoff-Nahoum et al., 2004, Slack et al., 2009, Tennant et al., 2008, Watnick and Jugder, 2020). Formicine ant species possess a highly acidic poison gland secretion containing formic acid as its main component (Lopez et al., 1993, Osman and Brander, 1961, Schmidt, 1986). Although the poison is presumably foremost used as a defensive weapon (Osman and Kloft, 1961), it is also distributed to the environment of these ants as an external immune defence trait (*sensu* Otti et al., 2014) to protect their offspring and the nest and to limit disease spread within the society (see references in Tragust, 2016, Brütsch et al., 2017, Pull et al., 2018). To this end, ants take up their poison from the acidopore, the opening of the poison gland at the gaster tip, into their mouth (Tragust et al., 2013) during a specialized behaviour existing only in a subset of ant families among all Hymenopterans (Basibuyuk and Quicke, 1999, Farish, 1972), termed acidopore grooming.

Here we first investigate whether the poison is also swallowed during acidopore grooming in *C. floridanus* and seven other formicine ant species from three genera in a comparative survey through measurement of pH levels in the crop and midgut lumen, experimental manipulation of acidopore access, and behavioural observations. In loss of acidopore and thus poison access during survival experiments and in *in vitro* and *in vivo* bacterial viability and growth experiments, we then investigate whether swallowing of the poison can serve gut microbial control and prevent bacterial pathogen infection analogous to acidic stomachs of higher vertebrates and acidic midgut regions in the fruit fly. Complementing these experiments, we also investigate whether acidopore access has the potential to limit pathogen transmission during trophallactic food exchange. Finally, we explore whether swallowing of the poison acts as a microbial filter that is permissible to gut colonization by bacteria from the family Acetobacteraceae.

## Results

### Swallowing of the poison gland secretion leads to acidic crops

Formicine ants naturally perform a specialized grooming behaviour, acidopore grooming, during which they take up their poison from the acidopore, the opening of the poison gland at the gaster tip, into their mouth (Basibuyuk and Quicke, 1999, Farish, 1972, Tragust et al., 2013). To reveal whether the poison is also swallowed during acidopore grooming, we monitored acidity levels in the crop lumen of the Florida carpenter ant *Camponotus floridanus* after feeding them 10% honey water (pH = 5). We found that after feeding the crop became increasingly acidic over time, reaching highly acidic values 48h after feeding (median pH = 2; 95% CI: 1.5-3.4), whilst renewed access to food after 48h raised the pH to levels recorded after the first feeding (Fig. 1a; LMM, LR-test, χ^2^ = 315.18, df = 3, *P* < 0.001; Westfall corrected post-hoc comparisons: 0+4h vs. 48h+4h: *P* = 0.317, all other comparisons: *P* < 0.001). This acidification after feeding was limited to the crop and did not extend to the midgut (Fig. 1 – figure supplement 1; pH-measurements at four points along the midgut 24h after access to 10% honey-water; mean ± se; midgut position 1 = 5.08 ± 0.18, midgut position 2 = 5.28 ± 0.17, midgut position 3 = 5.43 ± 0.16, midgut position 4 = 5.31 ± 0.19). Prevention of acidopore grooming in *C. floridanus* ants through immobilisation (see Materials and Methods) for 24h after feeding resulted in a significantly diminished acidity in the crop compared to non-prevented ants (Fig. 1b; LMM, LR-test, χ^2^ = 44.68, df = 1, *P* < 0.001). A similar, significantly diminished acidity in crop lumens was ubiquitously obtained in a comparative survey across seven formicine ant species (genera: *Camponotus*, *Lasius* and *Formica*) when ants were prevented from acidopore grooming through immobilisation compared to non-prevented ants (Fig. 1c; two-sided Wilcoxon rank sum tests, comparisons for all ant species: *P* ≤ 0.036). This indicates that the causative agent of crop acidity in formicine ants is not an internal source but an external one and that acidity of crop lumens is achieved through swallowing of the acidic poison during acidopore grooming.

**Fig. 1.**
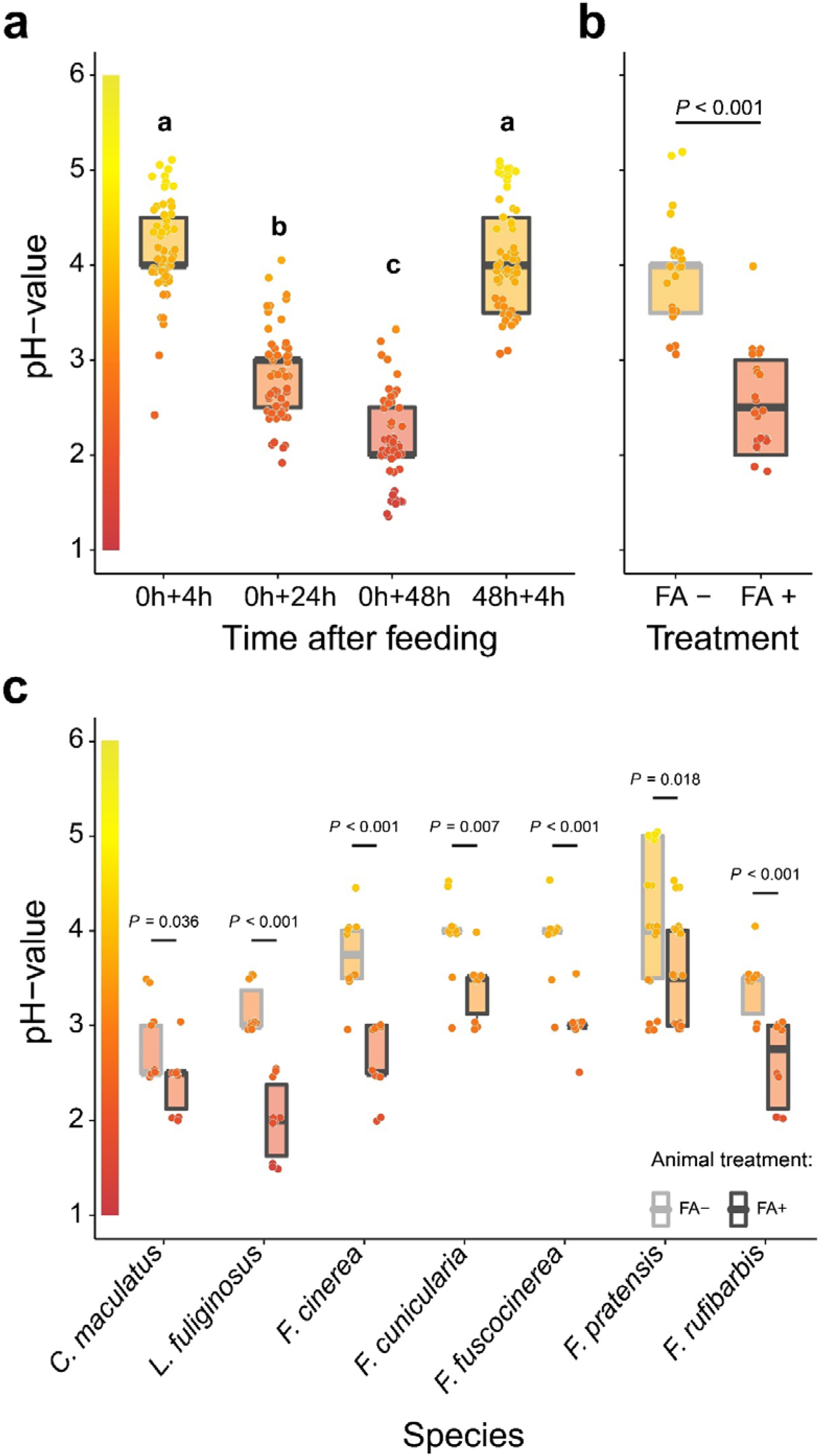
Acidification of formicine ant crop lumens through swallowing of acidic poison gland secretions. **a**, pH of crop lumens at 4h, 24h and 48h after feeding *C. floridanus* ants 10% honey water (pH = 5) at 0h and at 4h after re-feeding ants at 48h (LMM, LR-test, χ^2^ = 315.18, df = 3, *P* < 0.001, same letters indicate *P* = 0.317 and different letters indicate *P* < 0.001 in Westfall corrected post hoc comparisons). **b**, pH of crop lumens in *C. floridanus* ants that were either prevented to ingest the formic acid containing poison gland secretion (FA−) or not (FA+) for 24h after feeding (LMM, LR-test, χ^2^ = 44.68, df = 1, *P* < 0.001). **c**, pH-value of crop lumens 24h after feeding in seven formicine ant species that were either prevented to ingest the formic acid containing poison gland secretion (FA−) or not (FA+). Wilcoxon rank sum tests (two-sided). Lines and shaded boxes show median and interquartile range; points show all data. Colours in shaded rectangles near y-axis represent universal indicator pH colours. Colour filling of shaded boxes correspond to median pH colour of x-axis groups and colour filling of points correspond to universal indicator pH colours. Border of shaded boxes represents animal treatment (light grey: prevention of poison ingestion, FA−; dark grey: poison ingestion not prevented, FA+).

Although venomous animals often bear a cost of venom production and express behavioural adaptations to limit venom expenditure (Casewell et al., 2013), *C. floridanus* increases the frequency of acidopore grooming within the first 30 min. after ingesting food but also after ingesting water compared to unfed ants (Fig. 1 – figure supplement 2; GLMM, LR-test, χ^2^ = 33.526, df = 2, *P* <0.001; Westfall corrected post-hoc pairwise comparisons, water vs. 10% honey-water: *P* = 0.634, unfed vs water and unfed vs 10% honey-water: *P* < 0.001). This suggests that the behaviour of acidopore grooming is temporarily upregulated after fluid ingestion, irrespective of the fluid’s nutritional value. Moreover, pH levels were highly acidic in the crop of *C. floridanus* ants taken directly out of a satiated colony (Fig. 1 – figure supplement 3; major workers: median pH = 2, 95% CI: 2-3; minor workers: median = 3, CI: 2.5-3.6) and in worker cohorts that were satiated for three days and then starved for 24h before measurements (majors: pH = 2, 95% CI: 2-3; minors: median = 2, CI: 2-3). This indicates that the increased frequency of acidopore grooming after fluid ingestion and swallowing of the acidic poison serves to maintain an optimal, acidic baseline pH in the crop lumen of formicine ants following perturbation thereof through ingested fluids.

### Poison acidified crops can serve microbial control

Acidic stomachs of vertebrates and acidic midgut regions in the fruit fly *Drosophila melanogaster* can prevent infection by bacterial pathogens (Giannella et al., 1972, Howden and Hunt, 1987, Martinsen et al., 2005, Overend et al., 2016, Tennant et al., 2008). For swallowing of the poison to play a similar role in formicine ants and for the ensuing acidification to take effect after perturbation of the crop pH through ingested fluids, ingested fluids need to remain in the crop for a minimum time before being passed to the midgut. We therefore estimated food passage over time through the gut of *C. floridanus* ants with food containing fluorescent particles. We found that only a small amount of ingested food is passed from the crop to the midgut 2-4h after feeding, while thereafter food is steadily passed from the crop to the midgut until 18h after feeding (Fig. 2 – supplementary figure 1).

**Fig. 2.**
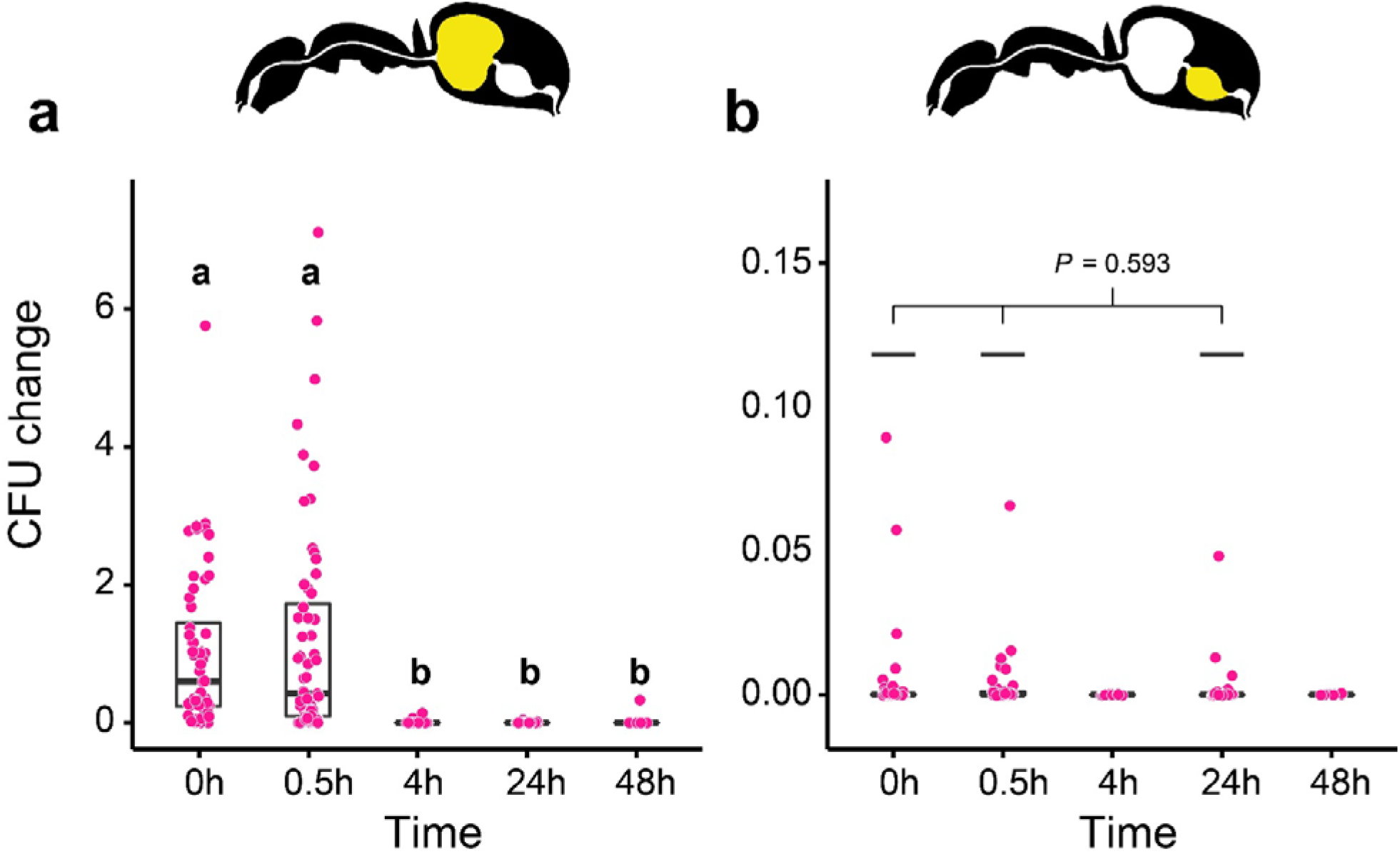
Viability of *S. marcescens* over time in the digestive tract of *C. floridanus*. Change in the number of colony forming units (CFUs) in the crop (**a**) and midgut (**b**) part of the digestive tract (yellow colour in insert) relative to the mean CFU-number at 0h in the crop (CFU change corresponds to single data CFU-value divided by mean CFU-value at 0h in the crop), 0h, 0.5h, 4h, 24h, and 48h after feeding *Camponotus floridanus* ants 10% honey water contaminated with *Serratia marcescens*. **a**, Change of *S. marcescens* in the crop (GLMM, LR-test, χ^2^ = 220.78, df = 4, *P* <0.001, same letters indicate *P* ≥ 0.623 and different letters indicate *P* < 0.001 in Westfall corrected post hoc comparisons). **b**, Change of *S. marcescens* in the midgut (GLMM, LR-test, χ^2^ = 1.044, df = 2, *P* = 0.593). Note that timepoints with zero bacterial growth in the midgut (4h and 48h) were excluded from the statistical model.

We then tested whether crop acidification after feeding could inhibit *Serratia marcescens*, an insect pathogenic bacterium (Grimont and Grimont, 2006), ingested together with food and prevent its passage from the crop to the midgut in *C. floridanus* ants. We measured this at two time points before (0.5h and 4h) and after (24h and 48h) main food passage from the crop to the midgut, with the time directly after food ingestion (0h) serving as a reference. When fed to *C. floridanus*, *S. marcescens* presence decreased sharply over time in the crop (Fig. 2a; GLMM, LR-test, χ^2^ = 220.78, df = 4, *P* < 0.001). The proportion of CFUs that we were able to retrieve from the crop relative to the mean at 0h in the crop diminished from 43% at 0.5h post-feeding (median, CI: 0-543%) to 0% at 4h (CI: 0-4%), 24h (CI: 0-1.8%), and 48h (CI: 0-18%) post-feeding. In addition, relative to the mean at 0h in the crop, *S. marcescens* could only be detected at extremely low numbers in the midgut (median 0%) at 0h (CI: 0-4%), 0.5h (CI: 0-1%) and 24h (CI: 0-1%) post-feeding and not at all at 4h and 48h post-feeding (Fig. 2b; GLMM, LR-test, χ^2^ = 1.044, df = 2, *P* = 0.593). This suggests that a lowering of the crop lumen pH through swallowing of the poison quickly and effectively reduces the viability of *S. marcescens* in the crop thus preventing further transport to the midgut.

An *in vitro* experiment on the ability of *S. marcescens* to withstand acidic conditions created with formic acid, the main component of the formicine poison gland secretion (Lopez et al., 1993, Osman and Brander, 1961, Schmidt, 1986), supports this interpretation. Incubation of *S. marcescens* for 2h in 10% honey water acidified with formic acid to pH 4 resulted in a significantly lower number of CFUs relative to pH 5 and in zero growth for incubations at pH-levels that were lower than 4 (Fig. 2 – figure supplement 2; GLM, LR-test, χ^2^ = 79.442, df = 1, *P* < 0.001). This *in vitro* experiment together with our *in vivo* experiment on the ability of *S. marcescens* to withstand acidic conditions in the crop of *C. floridanus*, indicates that poison acidified crops can serve microbial control in formicine ants.

### Acidopore access improves survival in the face of pathogen contaminated food

To test whether the ability to swallow the acidic poison can limit infection by oral pathogens, we prevented acidopore grooming through immobilisation in *C. floridanus* ants for 24h after feeding them once with 5μl of either honey water contaminated with *S. marcescens*, or non-contaminated honey water and monitored their survival thereafter without providing additional food. We found that acidopore access after pathogen ingestion significantly increased the survival probability of ants (Fig. 3; COXME, LR-test, χ^2^ = 20.95, df = 3, *P* = 0.0001). The survival of ants prevented from acidopore grooming and fed once with the pathogen contaminated food was significantly lower than that of non-prevented ants fed the same food source (Westfall corrected post-hoc comparisons: FA - | *Serratia* presence + vs. all other ant groups: *P* ≤ 0.027). Contrary, non-prevented ants fed once with the pathogen contaminated food source did not differ in survival to prevented and non-prevented ants fed the non-contaminated food source (Westfall corrected post-hoc comparisons: FA + | *Serratia* presence + vs. FA + | *Serratia* presence – and FA + | *Serratia* presence + vs. FA + | *Serratia* presence –: *P* ≥ 0.061 both). This indicates that acidopore access and thus swallowing of the poison provides a fitness benefit in terms of survival to formicine ants upon ingestion of pathogen contaminated food.

**Fig. 3.**
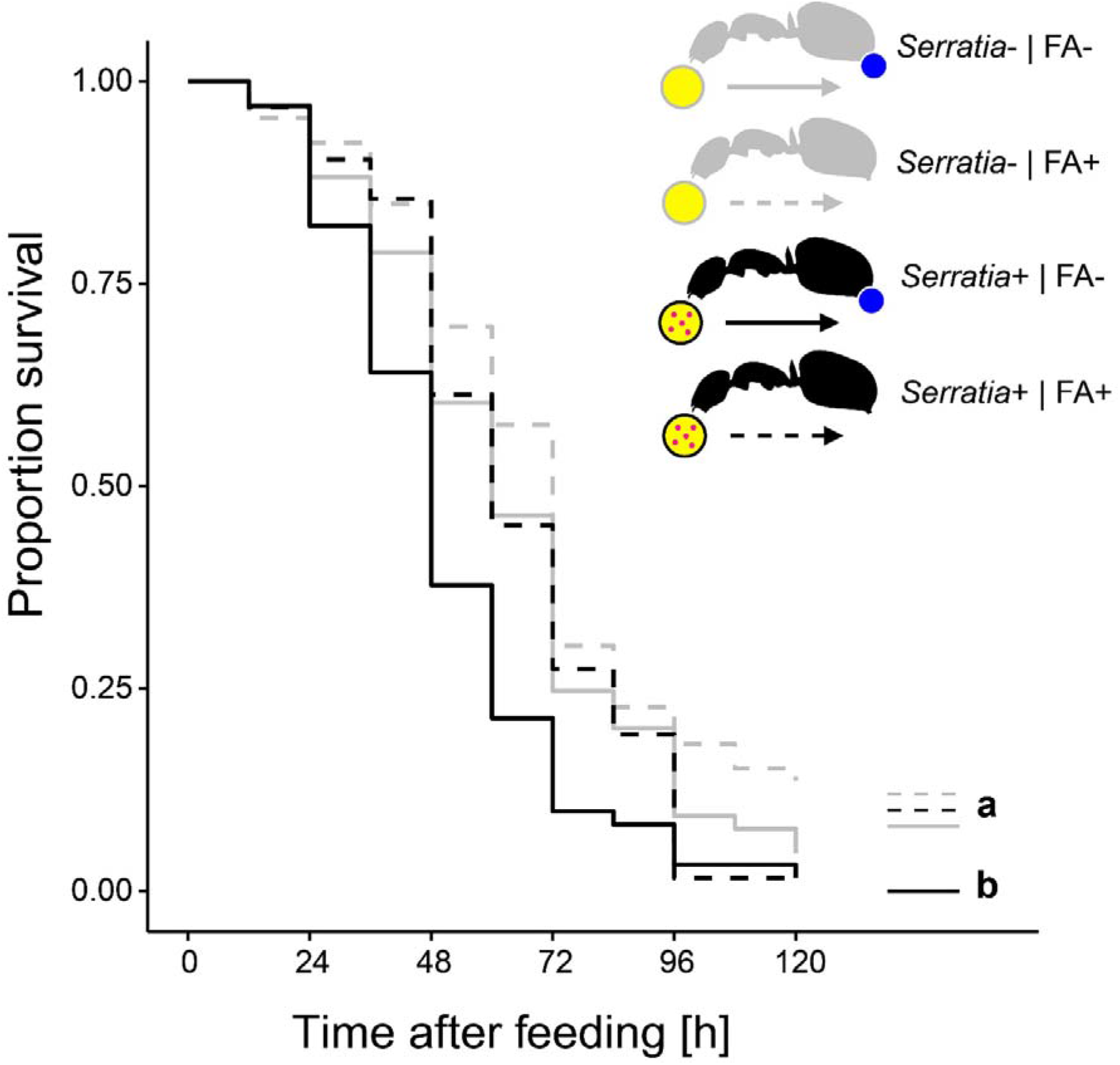
Survival after ingestion of pathogen contaminated food. Survival of individual *C. floridanus* ants that were either prevented to ingest the formic acid containing poison gland secretion (FA−; ant outlines with blue dot) or not (FA+) after feeding them once either honey water contaminated with *Serratia marcescens* (*Serratia*+, yellow circle with pink dots and black ant outlines) or non-contaminated honey water (*Serratia*-) without providing food thereafter (COXME, LR-test, χ^2^ = 20.95, df=3,*P* = 0.0001, same letters indicate *P* ≥ 0.061 and different letters indicate *P* ≤ 0.027 in Westfall corrected post hoc comparisons).

### Acidopore access can limit pathogen transmission

The ability to swallow the acidic poison may not only improve survival of formicine ants feeding directly on pathogen contaminated food but also of ants that share the contaminated food via trophallaxis. To test this, we created two types of donor-receiver ant pairs. Donor ants in both pairs were directly fed *S. marcescens* contaminated food every other day, while receiver ants obtained food only through trophallaxis from their respective donor ants. Receiver ants in both pairs were precluded from swallowing of the poison through blockage of their acidopore opening (see Materials and Methods), while donor ants were blocked in one pair but only sham blocked in the other pair. We found that the duration of trophallaxis between the two donor-receiver ant pairs during the first 30min. of the first feeding bout did not significantly differ (Fig. 4 – figure supplement 1; LMM, LR-test, χ^2^ = 1.23, df = 1, *P* = 0.268), indicating that trophallactic behaviour was not influenced through acidopore blockage in donor ants at the beginning of the experiment. Over the next 12 days we found that acidopore blockage *per se* had a significant negative effect on the survival of donor as well as receiver ants (Fig. 4; COXME, LR-test, χ^2^ = 66.68, df = 3, *P* < 0.001). Importantly however, although receiver ants that obtained food every other day from donors with the ability to swallow the poison died at a higher rate than their respective donor counterparts (hazard ratio: 1.81; Westfall corrected post-hoc comparison: *P* < 0.001) they were only half as likely to die compared to receiver ants that obtained pathogen contaminated food from blocked donors unable to swallow the poison (hazard ratio: 0.56; Westfall corrected post-hoc comparison: *P* < 0.001). This indicates that swallowing of the poison after feeding on pathogen contaminated food does not only improve survival of formicine ants directly feeding on pathogen contaminated food but also of ants that share the contaminated food via trophallaxis. Hence, swallowing of the poison and the ensuing crop acidity have the potential to limit oral disease transmission during food distribution within a formicine ant society.

**Fig. 4.**
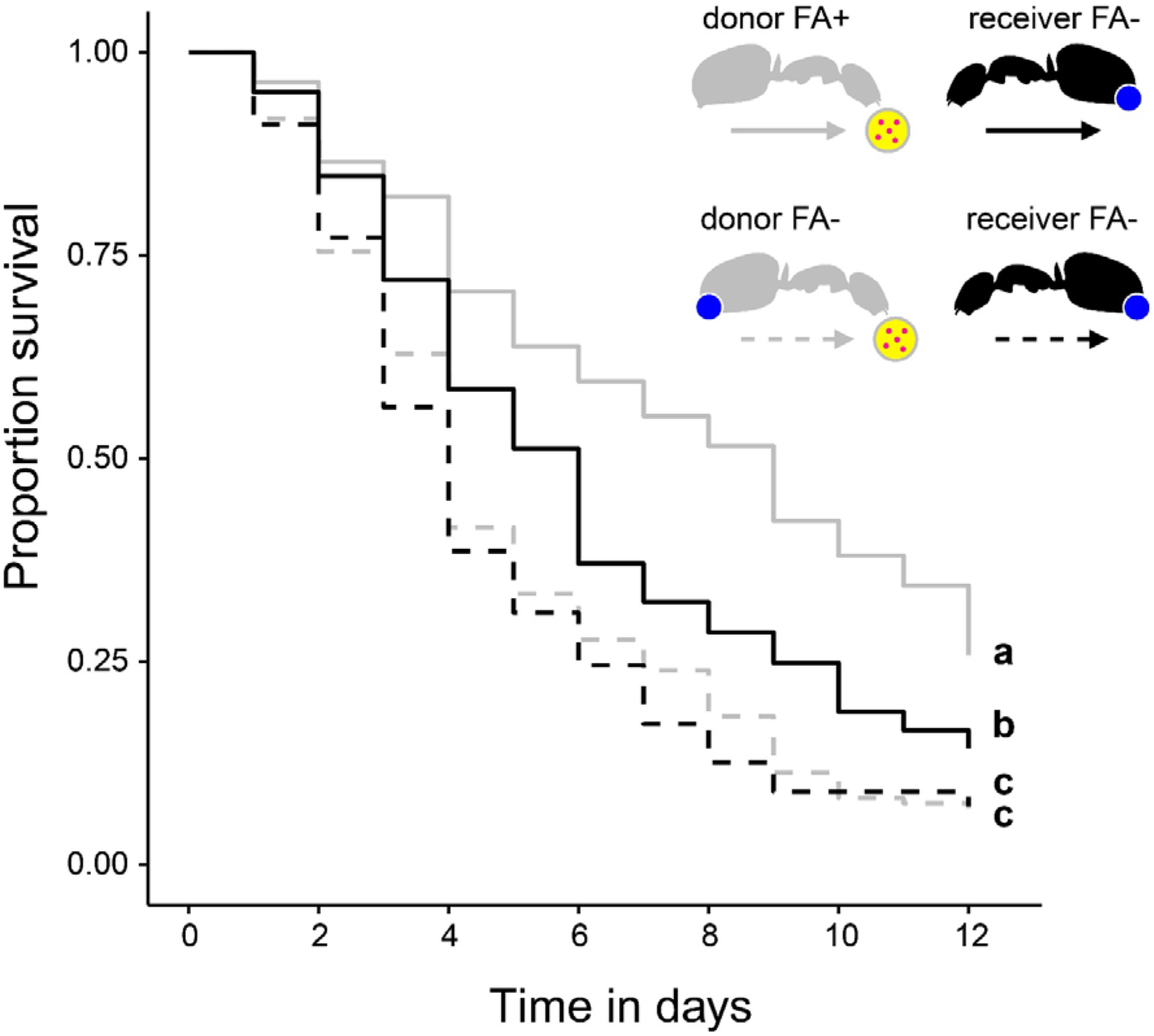
Survival after sharing pathogen contaminated food via trophallaxis. Survival of donor ants (light grey ant outlines) that were directly fed with pathogen contaminated food (yellow circle with pink dots in insert) every other day and were either prevented to ingest their formic acid containing poison gland secretion (FA−; ant outlines with blue dot) or not (FA+) and survival of receiver ants (black ant outlines) that received pathogen contaminated food only through trophallaxis with donor ants and were always prevented to ingest their formic acid containing poison gland secretion (FA−) (COXME, LR-test, χ^2^ = 66.68, df = 3,*P* < 0.001, same letters indicate *P* = 0.309 and different letters indicate *P* ≤ 0.002 in Westfall corrected post hoc comparisons).

### Poison acidified crops can serve as a chemical filter for microbial selection

In addition to pathogen control, poison acidified formicine ant crops might act as a chemical filter for gut associated microbial communities, similar to gut morphological structures that can act as mechanical filters in ants and other insects (Itoh et al., 2019, Lanan et al., 2016, Ohbayashi et al., 2015). To investigate the idea of a chemical filter, we tested the ability of the insect gut associated bacterium *Asaia* sp. (family Acetobacteraceae) (Crotti et al., 2009, Favia et al., 2007) to withstand acidic environments *in vitro* and *in vivo*. In contrast to *S. marcescens* (Fig. 2 – supplementary figure 2), *Asaia* sp. was not affected by an incubation for 2h in 10% honey water acidified with formic acid to a pH of 4 and was still able to grow when incubated at a pH of 3 in *in vitro* tests (Fig. 5 – figure supplement 1; GLM, overall LR-test χ^2^ = 21.179, df = 2, *P* < 0.001; Westfall corrected post hoc comparisons: pH = 5 vs. pH = 4: *P* = 0.234, all other comparisons: *P* < 0.001). Moreover, in *in vivo* tests, *Asaia* sp. only gradually diminished over time in the crop (Fig. 5a; GLMM; LR-test, χ^2^ = 124.01, df = 4, *P* < 0.001) with the proportion of CFUs that we were able to retrieve from the crop relative to the mean at 0h in the crop diminishing to only 34% (median, CI: 3-85%) and 2% (CI: 0-7%) at 4h and 24h post-feeding, respectively. At the same time, relative to the mean at 0h in the crop, *Asaia* sp. steadily increased in the midgut (Fig. 5b; GLMM; LR-test, χ^2^ = 59.94, df = 3, *P* < 0.001) from its initial absence at 0h post-feeding to 2% (median, CI: 0-5%) at 48h post-feeding. This indicates that a lowering of the crop lumen pH through swallowing of the acidic poison is permissible to the passage of *Asaia* sp.from the crop to the midgut. Together our *in vitro* and *in vivo* experiments with *S. marcescens* and *Asaia* sp. suggest that in formicine ants, poison acidified crops might act as a chemical filter that works selectively against the establishment of opportunistic and potentially harmful bacteria but allows entry and establishment of members of the bacterial family Acetobacteraceae. This view is also supported by results obtained *in vivo* for *E. coli*, a bacterium that is not a gut associate of insects (Blount, 2015) and which, similarly to *S. marcescens*, was not able to pass from the crop to the midgut in *C. floridanus* ants (Fig. 5 – figure supplement 2; crop: GLMM, LR-test, χ^2^ = 156.74, df = 4, *P* < 0.001; midgut: GLMM, LR-test, χ^2^ = 14.898, df = 3, *P* = 0.002).

**Fig. 5.**
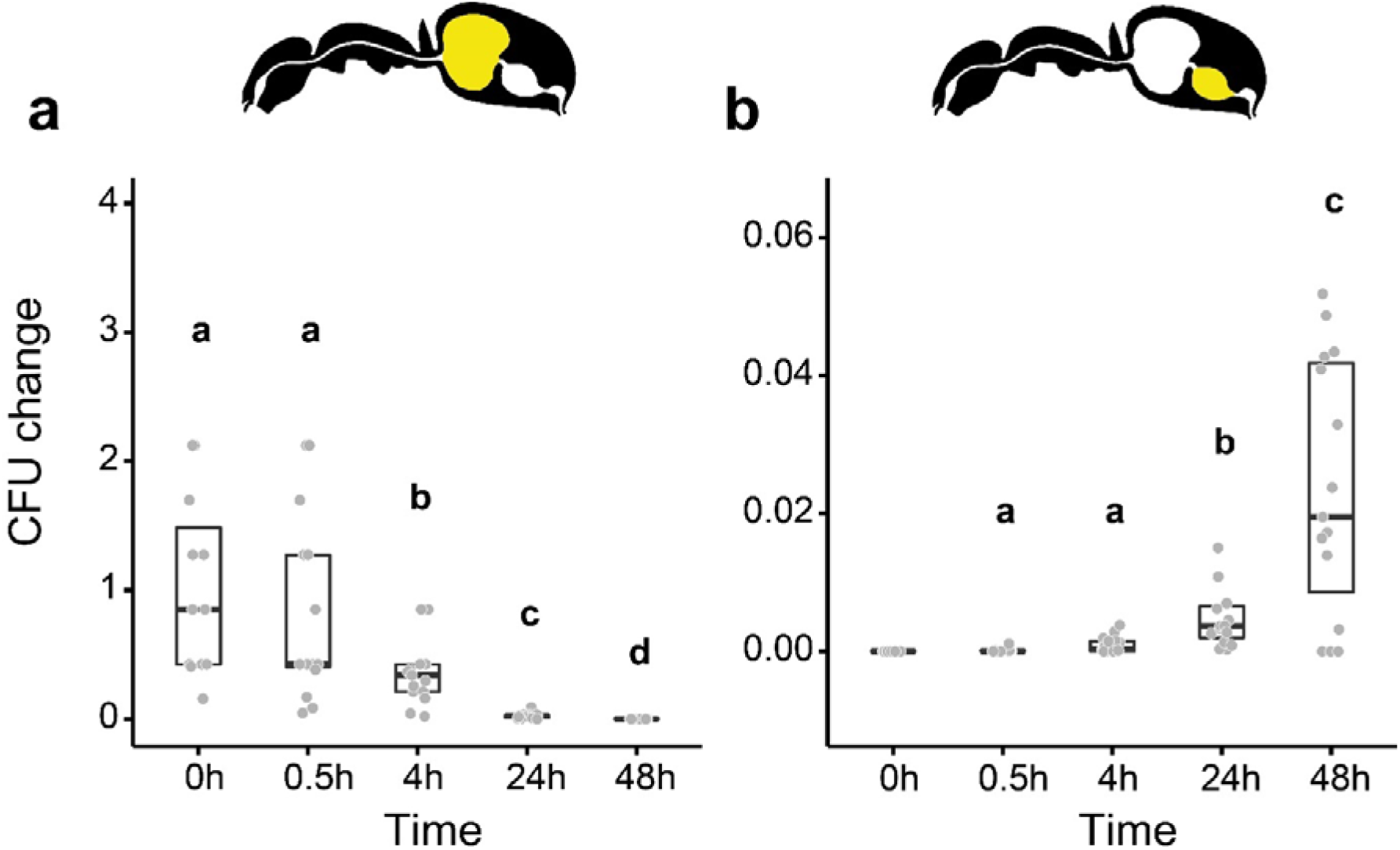
Viability of *Asaia* sp. over time in the digestive tract of *C. floridanus*. Change in the number of colony forming units (CFUs) in the crop (**a**) and midgut (**b**) part of the digestive tract (yellow colour in insert) relative to the mean CFU-number at 0h in the crop (CFU change corresponds to single data CFU-values divided by mean CFU-value at 0h in the crop), 0h, 0.5h, 4h, 24h, and 48h after feeding ants 10% honey water contaminated with *Asaia* sp. **a**, Change of *Asaia* sp. in the crop (GLMM; LR-test, χ^2^ = 124.01, df = 4,*P* < 0.001, same letters indicate *P* = 0.488 and different letters indicate *P* ≤ 0.013 in Westfall corrected post hoc comparisons). **b**, Change of *Asaia* sp. in the midgut (GLMM; LR-test, χ^2^ = 59.94, df = 3,*P* < 0.001, same letters indicate *P* = 0.116 and different letters indicate *P* ≤ 0.005 in Westfall corrected post hoc comparisons). Note that timepoints with zero bacterial growth in the midgut (0h) were excluded from the statistical model.

## Discussion

The results of our study indicate that formicine ants not only distribute their highly acidic, antimicrobial poison to their environment as an external immune defence trait (see references in Tragust, 2016, Brütsch et al., 2017, Pull et al., 2018), but also swallow the poison resulting in the creation of an acidic environment in their stomach, the crop (Fig. 1). This acidic environment can serve microbial control, limiting establishment of orally acquired, pathogenic microbes with ingested food (Fig. 2). In line with this, survival of formicine ants that fed pathogen contaminated food was improved if the ants were able to swallow their poison (Fig. 3). This fitness benefit also extended to ants that received pathogen contaminated food via trophallaxis from donor ants with the ability to swallow their poison (Fig. 4). Additionally, poison acidified crop lumens can act as chemical filters that selectively allow gut entry and establishment of members of the bacterial family Acetobacteraceae, a recurring part of the gut microbiota of formicine ants (Fig. 5) (Brown and Wernegreen, 2016, Chua et al., 2018, He et al., 2011, Ivens et al., 2018, Russell et al., 2017).

Instances of extreme pH conditions, especially alkaline, have been described in gastrointestinal tract compartments of several insects (Chapman, 2013, Holtof et al., 2019). Our study is however the first to report highly acidic conditions in insect crops and in animal stomachs in general apart from higher vertebrate species (Beasley et al., 2015, Koelz, 1992). Acidic conditions in the crop and other gut compartments of insects might be achieved in several ways. In principle, a digestive compartment with a certain pH can be generated through physiological mechanisms involving a transport-loop of acid-base equivalents across epithelia (Onken and Moffett, 2017). Insects could thus regulate the pH of their crop or of other gut compartments through active uptake and excretion of acid–base equivalents across the gut epithelium (Matthews, 2017). With few notable exceptions (Flower and Filshie, 1976, Miguel-Aliaga et al., 2018), the exact physiological mechanisms responsible for the creation of a gut lumen compartment with a certain pH are however often unknown (Harrison, 2001). Alternatively, insect gut associated microbes might contribute to the gut pH. Many bacteria produce short chain fatty acids and are thus able to acidify their environment (Ratzke et al., 2018, Ratzke and Gore, 2018). Members of the bacteria family Acetobacteraceae, and various Lactobacilli, gut microbial associates of Hymenoptera (McFrederick et al., 2013), release acetic acid as a waste product of their fermentative metabolism (Oude Elferink et al., 2001, Wolfe, 2005) and might thus influence pH conditions in the gut of their insect host. Finally, glandular secretions might contribute to the pH of the insect gut. In grasshoppers and crickets it has been suggested that salivary glands are responsible for slightly acidic (pH 5-6) crop lumens (Cooper and Vulcano, 1997, Harrison et al., 1992) and a recent study in the honeybee found that larval food has a pH of 4 due to the addition of fatty acids from a cephalic gland connected to the oral cavity (Muresan and Buttstedt, 2019). The oral cavity of ants is connected to several cephalic and thoracic glands (Emmert, 1968, Hölldobler and Wilson, 1990) that apart from lubricating compounds and digestive enzymes (Billen, 2009) can also produce acidic substances in some ants (Attygalle and Morgan, 1984, Morgan, 2008, Vander Meer, 2012). Physiological mechanisms, gut associated microbes or cephalic and thoracic glands connected to the oral cavity are however unlikely to be responsible for crop lumen acidity in our study, as immobilisation upon food ingestion severely impaired acidification of the crop in all formicine ants tested. This rather indicates that formicine ants use an external source for acidification but not an internal one.

Ants possess a huge diversity of exocrine glands all over their body (Billen, 2009, Billen and Sobotnik, 2015). Several of these glands produce acidic secretions (Attygalle and Morgan, 1984, Morgan, 2008, Vander Meer, 2012) and could thus serve as external sources for crop acidity. Most notable among these exocrine glands are the metapleural gland and the poison gland whose openings are located on the thorax and the gaster tip, respectively. Both produce acidic secretions in several ant species (Fernández-Marín et al., 2015, Lopez et al., 1993, Osman and Brander, 1961, Yek and Mueller, 2011) and are actively groomed with movements involving the mouth (Basibuyuk and Quicke, 1999, Farish, 1972, Fernández-Marín et al., 2009, Fernández-Marín et al., 2006, Tragust et al., 2013). Although immobilisation of ants in our experiments effectively prevented grooming of both glands, acidic metapleural gland secretions cannot serve as a source for crop acidity in the focal ant species of our study, *Camponotus floridanus*, and in the ant *Camponotus maculatus* in our comparative survey as ants of the genus *Camponotus* show, with few exceptions, an evolutionary loss of the metapleural gland (Yek and Mueller, 2011). In addition, in the formicine ant *Lasius neglectus*, grooming of the poison gland opening, the acidopore, results in the uptake of the poison into the mouth, while metapleural gland grooming was never observed (Tragust et al., 2013). Therefore, acidity of formicine ant crops in our study is likely the result of acidopore grooming and swallowing of the highly acidic poison, although our experiments cannot completely rule out the use of other sources.

Interestingly, although immobilisation of ants upon food ingestion led to a significantly impaired acidification of the crop in all formicine ants tested, we found a variable crop acidity in immobilised but also non-immobilised ants in our comparative survey. While this could indicate the use of acidic sources other than the poison gland, it could also indicate species specific differences in acidopore grooming, differences in the composition of the poison gland secretion or, as the ant crop is seldom completely empty (Greenwald et al., 2015, Greenwald et al., 2018) simply be the result of differences in crop filling status before feeding in our experimental setup. In *C. floridanus* we found that ants increased the frequency of acidopore grooming following ingestion of water or food. Together with the observed baseline level of acidity in the crop of around pH 2-3 under fully satiated and starved conditions, this indicates that *C. floridanus* ants strive to maintain an acidic, likely optimal pH in their crop through swallowing of their poison following perturbation of the crop pH after the ingestion of fluids. Although digestion is initiated in the crop, the midgut is the primary site of digestion in insects and the midgut epithelium plays a pivotal role in maintaining an optimal pH, as the pH is one of the most important regulators of digestive enzyme activity (Holtof et al., 2019, Terra and Ferreira, 1994). This might be the reason why the midgut pH of *C. floridanus* shows only slightly acidic levels (pH 5) after highly acidic levels in the crop; a change in pH that could be achieved via physiological mechanisms. While the variable acidity of crop lumens in our comparative survey will have to await further investigations, in *C. floridanus* ants, both, the increased frequency of acidopore grooming upon fluid ingestion and the change of pH to only slightly acidic levels in the midgut, the primary site of digestion, indicate that poison acidified crop lumens in formicine ants serve other functions than digestion, although an additional digestive function will have to be investigated.

Acidic fermentation of food is, for humans, a common practice which dates back centuries (Hutkins, 2019). Due to their antimicrobial activity (Cherrington et al., 1991, Hirshfield et al., 2003, Lund et al., 2014), the addition of organic acids to food represents the best practice to preserve food together with refrigeration (Theron and Rykers Lues, 2010). In *C. floridanus* ants ingested food largely remained in the crop for at least 2-4h before being passed to the midgut, which agrees with food passage times through the gastrointestinal tract of other ants (Cannon, 1998, Gößwald and Kloft, 1960, Gösswald and Kloft, 1960a, Howard and Tschinkel, 1981, Markin, 1970). This would enable swallowing of the poison and the ensuing crop acidity to take effect after perturbation of the crop pH through feeding. Acidic crop lumens in formicine ants might therefore play a role in the sanitation of ingested fluids similar to the ubiquitous, antimicrobials enabled sanitation of food in animals that provision food to their offspring or that store, cultivate, develop or live in food (Cardoza et al., 2006, Currie et al., 1999, Herzner et al., 2013, Herzner and Strohm, 2007, Joop et al., 2014, Milan et al., 2012, Mueller et al., 2005, Scott et al., 2008, Shukla et al., 2018a, Vander Wall, 1990, Vogel et al., 2017). A food sanitation function is supported by the rapid reduction of the insect bacterial pathogen *Serratia marcescens* in the crop of *C. floridanus* ants *in vivo* and the sensitivity of this bacterial pathogen to acidic conditions created with formic acid *in vitro*. The sensitivity of *S. marcescens* to acidic conditions created with formic acid *in vitro* likely even underestimates the antimicrobial effect of the formicine ant poison *in vivo*. Although formic acid is the main component of the formicine ant poison (Schmidt, 1986), the poison gland also contains acetic acid, hexadecanol, hexadecyl formate, hexadecyl acetat (Lopez et al., 1993) and a small fraction of unidentified peptides (Hermann and Blum, 1968, Osman and Brander, 1961). Additionally, the poison of formicine ants is usually expelled from the acidopore together with contents of the Dufour gland (Regnier and Wilson, 1968, Schoeters and Billen, 1996, but see Billen, 1982), which serve as wetting agents for the poison (Löfqvist, 1976, see also Kohl et al., 2001). In a previous study we found that formic acid alone could explain 70% of the antimicrobial activity of the poison against an entomopathogenic fungus, but that the combination of formic acid with other components of the poison gland and the Dufour gland could explain 94% (Tragust et al., 2013). In addition to the likely higher antimicrobial activity of the natural poison compared to the activity of formic acid alone, *in vivo* immune system effectors released into the gut lumen might contribute to the inability of *S. marcescens* to establish in the gastrointestinal tract of *C. floridanus.* Highly acidic stomachs in vertebrates and acidic midgut regions in the fruit fly *Drosophila melanogaster* serve together with immune system effectors microbial control and prevent infection by oral pathogens (Giannella et al., 1972, Howden and Hunt, 1987, Martinsen et al., 2005, Overend et al., 2016, Rakoff-Nahoum et al., 2004, Slack et al., 2009, Tennant et al., 2008, Watnick and Jugder, 2020). Future studies will need to investigate the contribution of immune system effectors released into the gut lumen to the rapid reduction of *S. marcescens* in the crop of *C. floridanus* and its inability to establish in the midgut.

Our study also found that access to the poison improved survival of *C. floridanus* ants after ingestion of food contaminated with the bacterial pathogen *S. marcescens*. Mortality of ants was generally high in this experimental setup irrespective of whether access to the poison was manipulated or not. This is unlikely the consequence of stress induced mortality through the method of immobilisation employed to prevent poison access in this experiment, as immobilised and non-immobilised ants did not differ in their survival upon ingestion of non-contaminated food. Rather, starvation following the one time feeding of contaminated and non-contaminated food likely increased mortality in all ants. In a similar experiment involving the formicine ant species *Formica exsecta*, oral exposure to *S. marcescens* contaminated food followed by starvation as well as starvation alone led to a high mortality of ants with no additive effects of pathogen exposure combined with starvation (Stucki et al., 2019). In addition to starvation, social isolation of *C. floridanus* ants might have increased mortality, as ants were kept individually in petri dishes. Social isolation has been shown to increase mortality in ants (Koto et al., 2015) and social isolation has been shown to reduce an individual’s capacity to fight infections in other group living animals (Kohlmeier et al., 2016). Despite the high mortality observed under these experimental conditions, our experiments on the ability of *S. marcescens* to withstand acidic conditions *in vitro* and *in vivo* and our survival experiment upon ingestion of *S. marcescens* contaminated food, suggest that swallowing of the poison and the ensuing crop acidity can indeed serve microbial control in formicine ants and improve survival in the face of pathogen contaminated food. In ants and other animals that lack an acidic poison, acids produced by other glands or acidic derivatives produced by defensive symbionts (Florez et al., 2015) or other environmental or gut associated bacteria, might provide functionally similar roles to the acidic poison in formicine anrts, as indicated in bees (Palmer-Young et al., 2018) and termites (Inagaki and Matsuura, 2018).

In addition to improve survival of ants that directly ingested pathogen contaminated food, the ability of donor ants to access their poison also improved survival of receiver ants without access to their poison when receiver ants shared pathogen contaminated food during trophallactic food exchange. Although our experiments on the ability of *S. marcescens* to withstand acidic conditions *in vivo* and *in vitro* indicate that this is likely due to fewer viable bacteria that are passed on from donor to receiver ants during trophallactic food exchange, it remains to be established whether this is indeed the case or whether this might be due to obtaining trophallactic fluids with antimicrobial activity. Antimicrobial activity of formicine ant trophallactic fluids has been described in previous studies (Hamilton et al., 2011, LeBoeuf et al., 2016). These studies linked the antimicrobial activity of trophallactic fluids to the presence of proteins related to cathepsin D, a lysosomal aspartic protease that can exhibit antibacterial effector activity and the proteolytic production of antimicrobial peptides (Ning et al., 2018). Our results however suggest a major role of swallowing the acidic poison to the antimicrobial activity of trophallactic fluids in formicine ants. Future studies will need to disentangle the relative contributions of crop acidity, proteins related to cathepsin D and, as previously pointed out, other immune effectors that are released into the insect gut to the antimicrobial activity of formicine ant trophallactic fluids.

The sensitivity of the bacterial pathogen *S. marcescens* to acidic conditions and the fitness benefit bestowed to ants with direct and indirect access to the poison after feeding on or receiving of pathogen contaminated food, might also indicate that swallowing of the poison and the ensuing crop acidity can act as an important barrier to oral disease spread within formicine ant societies. In the formicine ant *Formica polyctena*, food passage from the crop to the midgut is dependent upon whether food is directly eaten or is transferred via trophallaxis with only 20% of the honey water consumed directly seen in the midgut 4.5 h after feeding, while 77% of the honey water received during trophallaxis reaching the midgut 2.5 hours after feeding (Gösswald and Kloft, 1960a). Poison acidified crop lumens might therefore alleviate the cost of sharing pathogen contaminated food (Onchuru et al., 2018, Salem et al., 2015) and effectively counteract the generally increased risk of pathogen exposure and transmission associated with group-living (Alexander, 1974, Boomsma et al., 2005, Kappeler et al., 2015). On the other hand, trophallactic food exchange is a dynamic process that is dependent upon the functional role of the worker (Greenwald et al., 2015), food type (Buffin et al., 2011), and likely many other contexts. For example, it has repeatedly been reported that after a time of starvation food is distributed extremely quickly and efficiently via trophallaxis within an ant colony (Buffin et al., 2009, Gösswald and Kloft, 1960b, Markin, 1970, Sendova-Franks et al., 2010, Traniello, 1977, Wilson and Eisner, 1957). While this increases the threat of pathogen dissemination together with food, it has been suggested that dilution and mixing of food together with the existence of disposable ants specialised in food storage can mitigate the threat of pathogenic and noxious substances distributed together with food (Buffin et al., 2011, Sendova-Franks et al., 2010). Recently, it has been reported that ant social networks can plastically respond to the presence of a pathogen and that ants alter their contact network to contain the spread of a disease (Stroeymeyt et al., 2018). In our study, the first bout of trophallactic food exchange between donor and receiver ants was not affected by the manipulation of poison access. However, especially at later time points, potential changes in the amount of food transmitted cannot be excluded, as trophallaxis and feeding behaviour in general might depend on the infection status of ants engaging in trophallactic food exchange (Hite et al., 2020). In future studies, it will therefore be interesting to examine whether swallowing of the poison and the ensuing crop acidity truly acts as a barrier to oral disease spread within formicine ant societies, especially given the technical advances to track multiple individuals of a group simultaneously over time that have been made in recent years (Gernat et al., 2018, Greenwald et al., 2015, Imirzian et al., 2019, Stroeymeyt et al., 2018).

Our results on the ability of bacteria to withstand acidic environments *in vitro* and *in vivo* also provide evidence that swallowing of the acidic poison might not only serve microbial control but might also act as a chemical filter for microbes, working selectively against the establishment of pathogenic or opportunistic bacteria but allowing entry and establishment of species from the bacteria family Acetobacteraceae in *C. floridanus* ants. In ants, gut morphological structures, i.e. the infrabuccal pocket, an invagination of the hypopharynx in the oral cavity (Eisner and Happ, 1962), and the proventriculus, a valve that mechanically restricts passage of fluids from the crop to the midgut (Eisner and Wilson, 1952), can consecutively filter particles and microbes out of ingested food (Cannon, 1998, Glancey et al., 1981, Lanan et al., 2016, Little et al., 2006, Quinlan and Cherret, 1978). In other insects, gut morphological structures have been described to act as mechanical filters for gut associated microbes (Itoh et al., 2019, Ohbayashi et al., 2015). Given that acidic gut compartments in humans and fruit flies not only serve prevention of oral diseases but generally structure the gut microbiota of their hosts (Imhann et al., 2016, Overend et al., 2016), it is likely that poison acidified crop lumens can explain at least part of the recurrent presence of Acetobacteraceae in the gut of formicine ants and the otherwise reduced microbial diversity and abundance of gut associated microbes (Brown and Wernegreen, 2016, Brown and Wernegreen, 2019, Chua et al., 2018, Chua et al., 2020, Ivens et al., 2018, Russell et al., 2017).

Acetobacteraceae often thrive in sugar-rich environments (Mamlouk and Gullo, 2013). As many ants predominantly feed on sugars, e.g. extrafloral nectaries and aphid honeydew (Blüthgen et al., 2004, Blüthgen et al., 2000, Way, 1963), the presence of Acetobacteraceae in the gut of formicine ants could simply be a by-product of their dietary preferences (Anderson et al., 2012). On the other hand, some members of the Acetobacteraceae are pathogenic in humans and the fruit fly *Drosophila melanogaster* (Greenberg et al., 2006, Roh et al., 2008, Ryu et al., 2008) and most Acetobacteraceae produce metabolites that can potentially interfere with insect physiology and innate immunity (Chouaia et al., 2014). This may indicate that Acetobacteraceae found in formicine ants can act as pathogens. The ability of *Asaia* sp., a member of the Acetobacteraceae, to withstand acidic conditions *in vitro* and *in vivo* and to establish in the midgut of *C. floridanus* however indicate that host filtering (Mazel et al., 2018) might account for the pattern of recurring gut microbial communities in formicine ants. In addition, where studied in detail, Acetobacteraceae found associated with the gut of formicine ants show signs of genomic and metabolic adaptations to their host environment (Brown and Wernegreen, 2019, Chua et al., 2020), indicating coevolution and potentially also mutual benefit, though this has not formally been established (see also Mushegian and Ebert, 2016). The creation of an acidic pH crop environment in formicine ants that is easier to endure if colonizing microbes are mutualists agrees with the theoretical concept of screening, as opposed to signalling, as a means of partner choice in cross-kingdom mutualisms (Archetti et al., 2011a, Archetti et al., 2011b, Biedermann and Kaltenpoth, 2014, Scheuring and Yu, 2012). Signalling is a well-known mechanism of partner selection in the context of mate choice (Grafen, 1990, Zahavi, 1975) or plant-microbe mutualisms (Clear and Horn, 2019, Rebolleda-Gomez and Wood, 2019) and signals play an important role in the cross talk between the immune system and host associated microbes (Allaire et al., 2018, Fischbach and Segre, 2016, Moura-Alves et al., 2019, Villena et al., 2018). If the signal has a cost, only good quality individuals will find it profitable to advertise their quality, and therefore the signal will be honest. Contrary to signalling, where costly information is displayed to partners, in screening a costly environment is imposed on partners that excludes all but high-quality ones. Partner choice in a number of cross-kingdom mutualisms is readily explained by screening (see examples in Archetti et al., 2011a, Archetti et al., 2011b, Biedermann and Kaltenpoth, 2014, Scheuring and Yu, 2012) but experimental evidence is so far limited in insect-microbe associations (Innocent et al., 2018, Itoh et al., 2019, Ranger et al., 2018). Although our experiments can only indicate screening as a means of partner choice in formicine ants, the results of our study would provide support for the prediction that screening is more likely to evolve from a host’s defence trait against parasites (Archetti et al., 2011a, Archetti et al., 2011b), i.e. the highly acidic poison that creates a selective environment for microbes.

## Conclusion

Overall, our study provides evidence that swallowing of the formic acid containing poison gland secretion and the ensuing acidity in the crop lumen acts as a chemical filter for control and selection of microbes ingested with food, protecting formicine ants from food borne bacterial pathogens and structuring their gut associated microbial communities. Only recently, studies have started to acknowledge that antimicrobials in the environment of an organism have the potential dual role of microbial selection and control (Brinker et al., 2019b, Duarte et al., 2018, Innocent et al., 2018, Shukla et al., 2018a, Shukla et al., 2018b, Strohm et al., 2019, Tragust et al., 2020). Antimicrobials used as external immune defence traits (Otti et al., 2014) may generally not only serve pathogen protection and microbial control but may act as microbial filters to manage host associated microbes, be it in food or the environment, and thus contribute to a host’s ecological and evolutionary success. In the case of social species, externally acting antimicrobials might contribute to the ecological and evolutionary success of this group of insects by alleviating the increased risk of pathogen exposure and transmission associated with group living but allowing the acquisition and transmission of microbial mutualists.

## Methods

### Ant species and maintenance

Colonies of the carpenter ant *Camponotus floridanus* were collected in 2001 and 2003 in Florida, USA, housed in Fluon® (Whitford GmbH, Diez, Germany) coated plastic containers with plaster ground and maintained at a constant temperature of 25°C with 70% humidity and a 12h/12h light/dark cycle. They were given water *ad libitum* and were fed two times per week with honey water (1:1 tap water and commercial quality honey), cockroaches (*Blaptica dubia*) and an artificial diet (Bhatkar and Whitcomb, 1971). For comparison, workers of one other *Camponotus* species (*Camponotus maculatus*), collected close to Kibale Forest, Uganda, in 2003 and housed under identical conditions as *Camponotus floridanus* were used. Additionally, six other formicine ant species, one *Lasius* and five *Formica* species (*Lasius fuliginosus, Formica cinerea, Formica cunicularia, Formica fuscocinerea, Formica pratensis* and *Formica rufibarbis*) were collected in Bayreuth, Germany in 2012 and 2018 and kept for approximately two weeks prior experimental use at 20°C, 70% humidity and a 14h/10h light/dark cycle. Except otherwise noted only the small worker caste (“minors”) of *Camponotus* species was used.

### Acidity of the crop lumen and pH measurements

To determine whether formicine ants swallow their poison after feeding, we tracked changes in pH-levels of the crop lumen in *C. floridanus* ants over time. Before use in experimental settings, cohorts of ~100 ants were taken out of their natal colony (n = 6 colonies) into small plastic containers lined with Fluon® and starved for 24-48h. Thereafter, ants were put singly into small petri dishes (Ø 55 mm) with damp filter paper covered bottom, given access to a droplet of 10% honey water (w/v) for 2h before removing the food source and measuring the pH of the crop lumen after another 2h (group 0+4h: n = 60 workers), after 24h (group 0+24h: n = 59 workers) or 48h (group 0+48h: n = 52 workers). To assess the effect of renewed feeding, a separate group of *C. floridanus* ants were given access to 10% honey water 48h after the first feeding for 2h prior to measuring the pH of their crop lumen after another 2h (group 48h+4h: n = 60 workers). To measure the pH, ants were first cold anesthetized on ice, then their gaster was cut off with a fine dissection scissor directly behind the petiole and leaking crop content (1-3μl) collected with a capillary (5μl Disposable Micro Pipettes, Blaubrand intraMARK, Brand, Wertheim). The collected crop content was then emptied on a pH sensitive paper to assess the pH (Hartenstein, Unitest pH 1-11). This method of collecting crop content will invariably result in some mixing of crop lumen content with haemolymph. As the pH of the insect haemolymph ranges from only slightly acidic (pH ≥ 6.5) to near-neutral or slightly alkaline (pH ≤ 8.2) (Harrison, 2001, Matthews, 2017), this might have biased the results of our pH measurements to slightly higher pH values. As a reference point for food pH, we also measured the pH of 10% honey water on pH sensitive paper, which gave invariably pH = 5.

In addition, we measured the pH in the crop lumen and at four points in the lumen along the midgut (1^st^ measurement directly behind proventriculus to 4^th^ measurement one mm apical from insertion point of the Malpighian tubules) of *C. floridanus* workers that were fed 24 h prior to measurements with 10% honey-water. For these measurements worker guts were dissected as a whole and pH was measured in the crop (n = 2 workers from two colonies) and along the midgut (all midgut points n = 10, except point four with n = 9 workers from four different colonies) with a needle-shaped microelectrode (UNISENSE pH-meter; microelectrode with needle tip of 20μm diameter).

In formicine ants, oral uptake of the poison into the mouth is performed via acidopore grooming (Tragust et al., 2013). During this behavior ants bend their gaster forward between the legs and the head down to meet the acidopore, the opening of the poison gland, at the gaster tip (Basibuyuk and Quicke, 1999, Farish, 1972). In an additional experiment we therefore compared the crop lumen pH of *C. floridanus* workers from four different colonies that were either prevented to reach their acidopore (FA-ants) or could reach their acidopore freely (FA+ ants). To do this, we again allowed single ants access to 10% honey water for 2h after a starvation period, before cold anesthetizing them briefly on ice and immobilizing FA-ants (n = 22 workers) in a pipetting tip, while FA+ ants (n = 23 workers) remained un-manipulated. After 24h we measured the pH of the crop lumen as before.

To investigate whether swallowing of the acidic poison is widespread among formicine ants, the latter experiment was repeated for six additional formicine ant species (FA-ants: n = 10 workers except for *Formica pratensis* with n = 21; FA+ ants: n = 10 workers except for *Formica pratensis* with n=20; all ants: n = 1 colony) in the same fashion as described before with the exception that apart from *Formica pratensis* the crop lumen was collected through the mouth by gently pressing the ants’ gaster. Crop lumen of *Formica pratensis* ants was collected in the same fashion as crop lumen of *C. floridanus* ants.

To investigate whether the type of fluid and its nutritional value have an influence on the frequency of acidopore grooming in *C. floridanus*, the following experiment was performed. Cohorts of ~100 ants were taken out of their natal colony (n = 6 colonies) into small plastic containers and starved for 24-48h. Thereafter, ants were again put singly into small petri dishes (Ø 55 mm) and given access to either a 3μl droplet of 10% honey water (w/v, n = 126 ants, treatment: honey-water fed), a 3μl droplet of tap water (n = 128, water-fed) or to no fluid (n = 125, unfed). After acclimatization (unfed ants) or after swallowing of the fluid (honey-water and water-fed ants, both 1-2min.), all ants were filmed for the next 30min. (Logitech webcam c910). These videos were then analyzed for the frequency of acidopore grooming.

Finally, we measured the pH in the crop lumen of *C. floridanus* ants (n = 3 colonies) under satiated and starved conditions to estimate a baseline level of acidity in the crop. For this, ants taken out of satiated, twice per week fed colonies on the day of feeding were compared to ants that were maintained in cohorts of ~100 individuals for three days with access to 10% honey-water and then starved for 24h before measuring the pH in their crop (n = 10 major and 10 minor workers per colony and condition). The pH in the crop lumen was measured as described before by briefly cold anesthetizing ants an ice, collecting the crop content through the mouth by gently pressing the ants’ gaster and then emptying it on a pH sensitive paper (Hartenstein, Unitest pH 1-11).

### Bacterial strains and culture

As model entomopathogenic bacterium *Serratia marcescens* DSM12481 (DSMZ Braunschweig, Germany) was used. This bacterium is pathogenic in a range of insects (Grimont and Grimont, 2006) and has been detected in formicine ants, i.e. *Anoplolepis gracilipes* (Cooling et al., 2018) and *Camponotus floridanus* (Ratzka et al., 2011). While often non-lethal within the digestive tract, *S. marcescens* can cross the insect gut wall (Mirabito and Rosengaus, 2016, Nehme et al., 2007) and is highly virulent upon entry into the hemocoel (Flyg et al., 1980), not least due to the production of bacterial toxins (Hertle, 2005). As a model bacterial gut-associate of ants *Asaia* sp. strain SF2.1 (Favia et al., 2007), was used*. Asaia* sp. belongs to the family Acetobacteraceae, members of which often thrive in sugar-rich environments (Mamlouk and Gullo, 2013), such as honey-dew that ants like *C. floridanus* predominantly feed on. *Asaia* sp. is capable of cross-colonizing insects of phylogenetically distant genera and orders (Crotti et al., 2009, Favia et al., 2007) and can be a component of the gut associated microbial community of formicine and other ants (Chua et al., 2018, Kautz et al., 2013a, Kautz et al., 2013b). In addition to *S. marcescens* and *Asaia sp.*, *Escherichia coli* DSM6897 (DSMZ Braunschweig, Germany) was used as a model opportunistic bacterium that is not a gut-associate of insects. *E. coli* bacteria are a principal constituent of mammalian gut associated microbial communities but are commonly also found in the environment (Blount, 2015).

Bacterial stocks of *S. marcescens, Asaia* sp., and *E. coli* were kept in 25% glycerol at −80°C until use. For use, bacteria were plated on agar plates (LB-medium: 10g Tryptone, 5g Yeast extract, 20g Agar in 1L MilliQ-water, and GLY-medium: 25g Gycerol, 10g Yeast extract, 20g Agar in 1L MilliQ-water with pH adjusted to 5.0, for *S. marcescens/E. coli* and *Asaia* sp. respectively), single colony forming units (CFUs) were picked after 24h (*S. marcescens/E. coli*) or 48h (*Asaia* sp.) of growth at 30°C and transferred to 5ml liquid medium (LB-medium and GLY-medium minus agar for *S. marcescens/E. coli* and *Asaia* sp. respectively) for an overnight culture (24h) at 30°C. The overnight culture was then pelleted by centrifugation at 3000g, the medium discarded and resolved in 10% (w/v) honey water to the respective working concentration for the experiments. The concentration of a typical overnight culture was determined for *S. marcescens* and *Asaia* sp. by plating part of the overnight culture on agar plates and counting CFUs after 24h or 48h of growth at 30°C, for *S. marcescens* and *Asaia* sp. respectively. This yielded a concentration of 1.865 * 10^9^ ± 5.63 * 10^7^ (mean ± sd) bacteria per ml for *S. marcescens* and 5.13 * 10^8^ ± 8.48 * 10^6^ (mean ± sd) bacteria for *Asaia* sp.

### Survival experiments

In a first survival experiment we tested whether the ability to perform acidopore grooming within the first 24h after ingestion of pathogen contaminated food provides a survival benefit for individual *C. floridanus* ants. Ants from eight colonies were starved for 24-48h before start of the experiment, as described before, and then workers put singly in small petri dishes were either given access to 5μl of *S. marcescens* contaminated 10% honey water (9.33 * 10^9^ bacteria/ml; *Serratia+* ants: n = 127) or uncontaminated 10% honey water (*Serratia-* ants: n = 135) for 2 min. Thereafter, all ants were cold anaesthetized and approximately half of the *Serratia+* and the *Serratia-* ants (n = 65 and n = 69, respectively) immobilized in a pipetting tip, thus preventing acidopore grooming (FA-ants: n = 134) while the other half remained fully mobile (FA+ ants: n = 128). After 24h, FA-ants were freed from the pipetting tip to minimize stress. Mortality of the ants was monitored over 5 days (120h) every 12h providing no additional food, except the one time feeding of 5μl contaminated or uncontaminated honey water at the start of the experiment. We chose to provide no additional food after the one time feeding at the beginning of the experiment, as an altered feeding behavior, i.e. illness induced anorexia with known positive or negative effects on survival (Hite et al., 2020), might otherwise have influenced our results.

In an additional survival experiment, we investigated whether the ability to acidify the crop lumen has the potential to limit oral disease transmission during trophallactic food transfer. To this end, *C. floridanus* ants from seven colonies were again starved, divided randomly in two groups (donor and receiver ants, each n = 322) and their gaster marked with one of two colours (Edding 751). Additionally, to prevent uptake of the poison, the acidopore opening of all receiver ants (receiver FA−) and half of the donor ants (donor FA−) was sealed with superglue, while the other half of the donor ants were sham treated (donor FA+) with a droplet of superglue on their gaster (Tragust et al., 2013). We then paired these ants into two different donor-receiver ant pairs. Pairs with both donor and receiver ants having their acidopore sealed (donor FA−| receiver FA−) and pairs with only receiver ants having their acidopore sealed (donor FA+ | receiver FA−). Six hours after pairing, donor ants from both pairs were isolated and given access to 5μl of *S. marcescens* contaminated 10% honey water (1.865 * 10^9^ bacteria/ml) for 12h. Thereafter donor ants were again paired with the respective receiver ants for 12 h and all pairs filmed for the first 30min. (Logitech webcam c910). These videos were then analyzed for the duration of trophallaxis events donor-receiver ant pairs engaged in during the first bout of trophallactic food exchange. After this first feeding round, donor ants were fed in the same fashion, i.e. isolation for 12h with access to *S. marcescens* contaminated 10% honey water, every 48h, while they were maintained with the respective receiver ants for the rest of the time. This experimental design ensured that receiver ants were fed only through the respective donor ants with pathogen contaminated food. Survival of both, donor and receiver ants, was monitored daily for a total of 12 days.

### Bacterial viability and growth assays

We tested the ability of *S. marcescens* and *Asaia* sp. to withstand acidic environments *in vitro*, as well as their ability and the ability of *E. coli* to pass from the crop to the midgut *in vivo* when ingested together with food. In ants, gut morphological structures, i.e. the infrabuccal pocket, an invagination of the hypopharynx in the oral cavity (Eisner and Happ, 1962), and the proventriculus, a valve that mechanically restricts passage of fluids from the crop to the midgut (Eisner and Wilson, 1952), consecutively filter solid particles down to 2μm (Lanan et al., 2016) which would allow *S. marcescens* (Ø: 0.5-0.8μm, length: 0.9-2μm Grimont and Grimont, 2006), *Asaia* sp. (Ø: 0.4-1μm, length: 0.8-2.5μm Komagata et al., 2014), and *E. coli* (length: 1μm, width: 0.35μm, Blount, 2015) to pass. For the *in vitro* tests we incubated a diluted bacterial overnight culture (10^5^ and 10^4^ CFU/ml for *S. marcescens* and *Asaia* sp., respectively) in 10% honey water (pH = 5) and in 10% honey water acidified with commercial formic acid to a pH of 4, 3 or 2 for 2h at room temperature (*S. marcescens*: n = 15 for all pH-levels, except pH = 4 with n = 13; *Asaia* sp.: n = 10). Then we plated 100μl of the bacterial solutions on agar-medium (LB-medium and GLY-medium for *S. marcescens* and *Asaia* sp., respectively) and incubated them at 30°C for 24h (*S. marcescens*) or 48h (*Asaia* sp.) before counting the number of formed CFUs. For the *in vivo* tests *C. floridanus* ants from five (*Asaia* sp.), four (*E. coli*) or from six colonies (*S. marcescens*) were starved as before and then individually given access to 5μl of bacteria contaminated 10% honey water (*Asaia* sp. and *E. coli*: 1 * 10^7^ CFU/ml, *S. marcescens*: 1 * 10^6^ CFU/ml) for 2 min. To assess the number of CFUs in the digestive tract, i.e. the crop and the midgut, ants were dissected either directly after feeding (0h; *S. marcescens*: n = 60 workers; *Asaia* sp. and *E. coli*: n = 15 each), or at 0.5h (*S. marcescens*: n = 60; *Asaia* sp. and *E. coli*: n = 15 each), 4h (*S. marcescens*: n = 60; *Asaia* sp. and *E. coli*: n = 15 each), 24h (*S. marcescens*: n = 53; *Asaia* sp. and *E. coli*: n = 15 each) or 48h (*S. marcescens*: n = 19; *Asaia* sp. and *E. coli*: n = 15 each) after feeding. For dissection, ants were cold anesthetized, the gaster opened and the whole gut detached. The crop and the midgut were then separated from the digestive tract, placed in a reaction tube, mechanically crushed with a sterile pestle and dissolved in 100μl (*Asaia* sp. and *E. coli*) or 150μl (*S. marcescens*) phosphate buffered saline (PBS-buffer: 8.74g NaCl, 1.78g Na_2_HPO_4_,2H_2_O in 1L MilliQ-water adjusted to a pH of 6.5). The resulting solutions were then thoroughly mixed, 100μl streaked on agar-medium (LB-medium and GLY-medium for *S. marcescens*/*E.coli* and *Asaia* sp., respectively) and incubated at 30°C for 24h (*S. marcescens* and *E. coli*) or 48h (*Asaia* sp.), before counting the number of formed CFUs. No other bacteria (e.g. resident microbes) were apparent in terms of a different CFU morphology on the agar plates which agrees with the very low number of cultivable resident bacteria present in the midgut of *C. floridanus* (Stoll and Gross, unpublished results). This methodology cannot completely exclude that resident *S. marcescens* or species of Acetobacteraceae might have biased our count data by adding a background level of CFUs at all timepoints or by adding random outlier CFUs at specific timepoints. Both, background level CFU numbers and random outlier CFUs should however not influence observed patterns over time. The timepoints of 0h, 0.5h, 4h, 24h and 48h in *in vivo* bacterial growth assays were chosen according to literature describing passage of food from the crop to the midgut within 3-6h after food consumption in ants (Cannon, 1998, Gösswald and Kloft, 1960b, Gösswald and Kloft, 1960a, Howard and Tschinkel, 1981, Markin, 1970). They should thus be representative of two time points before food passage from the crop to the midgut (0.5h and 4h) and two time points after food passage from the crop to the midgut (24h and 48h) together with the reference timepoint (0h).

### Food passage experiment

To estimate food passage from the crop to the midgut and hindgut of *C. floridanus* after feeding we performed the following experiment. We again took a cohort of ~100 workers out of one natal colony of *C. floridanus*, starved them for 24h and then offered them 200 μl of a 1:1 honey-water mix with 50mg of polymethylmethacrylate (PMMA, aka acrylic glass) particles (size ≤ 40 μm). Thereafter, we dissected the digestive tract of three major and three minor workers at each of the timepoints 2h, 4h, 6h, 8h, 12h, 14h, 16h, 18h, 24h, and 48h after feeding and placed each under a microscope (Leica DM 2000 LED) to detect and count the number of particles via fluorescence in the crop, the midgut and the hindgut.

### Statistical analyses

All statistical analyses were performed with the R statistical programming language (version 3.6.1, R Core Team, 2019). All (zero-inflated) General(ized) linear and mixed models and Cox mixed-effects models were compared to null (intercept only) or reduced models (for those with multiple predictors) using Likelihood Ratio (LR) tests to assess the significance of predictors. Pairwise comparisons between factor levels of a significant predictor were performed using pairwise post-hoc tests adjusting the family-wise error rate according to the method of Westfall (package “multcomp”, Bretz et al., 2011). We checked necessary model assumptions of (zero-inflated) General(ised) linear and mixed models using model diagnostic tests and plots implemented in the package “DHARMa” (Hartig, 2019). Acidity of the crop lumen (log transformed pH to normalize data) and midgut lumen in *C. floridanus* was analyzed using linear mixed models (LMM, package “lme4”, Bates et al., 2015) including time since feeding (four levels: 0+4h, 0+24h, 0+48h, 48h+4h; Fig. 1a), ant manipulation (two levels: FA+ and FA−, i.e. ants with and without acidopore access; Fig. 1b) or digestive tract part (four levels: crop, midgut position 1, midgut position 2, midgut position 3, midgut position 4; Fig. 1 – figure supplement 1) as predictors and natal colony as a random effect. Due to non-normality and heteroscedasticity, the acidity of the crop lumen in the seven formicine ant species other than *C. floridanus* (Fig. 1c) was analysed using per species Wilcoxon Rank Sum tests with ant manipulation (FA+ and FA−) as predictor. The frequency of acidopore grooming in *C. floridanus* upon feeding different types of fluids was analyzed using Generalized linear mixed models (GLMM, package “lme4”, Bates et al., 2015) with negative binomial errors and type of fluid (three levels: unfed, water-fed or 10% honey water fed) as predictor and natal colony as random effect (Fig. 1 – figure supplement 2).

Survival data were analysed with Cox mixed effects models (COXME, package “coxme”, Therneau, 2019). For the survival of individual ants (Fig. 3), ant treatment (four levels: *Serratia-* | FA−, *Serratia-* | FA+, *Serratia*+ | FA−, *Serratia+* | FA+) was added as a predictor and the three “blocks” in which the experiment was run and the colony ants originated from, were included as two random intercept effects. For the survival of donor-receiver ant pairs (Fig. 4), ant treatment (four levels: donor FA+, donor FA−, receiver FA+, receiver FA−) was included as a predictor and the three “blocks” in which the experiment was run, the colony ants originated from, and petri dish in which donor and receiver ants were paired, were included as three random intercept effects. Survival of receiver ants was right censored if the corresponding donor ant died at the next feeding bout (right censoring of both donor and receiver ants in one pair upon death of one of the ants yielded statistically the same result: COXME, overall LR χ^2^ = 60.202, df = 3, P < 0.001; post-hoc comparisons: receiver FA-vs donor FA−: *P* = 0.388, all other comparisons: P < 0.001). The duration of trophallaxis events (square-root transformed to normalize data) between donor-receiver ant pairs was analysed using a linear mixed model with ant pair type (two levels: donor FA+ | receiver FA- and donor FA−| receiver FA−) as predictor and the three “blocks”, in which the experiment was run and the colony ants originated from as random effect (Fig. 4 – figure supplement 1).

Bacterial growth *in vitro* was analysed separately for *S. marcescens* and *Asaia* sp. using Generalized linear models (GLM) with negative binomial errors and pH as predictor, excluding pH levels with zero bacterial growth due to complete data separation (Fig. 2 – supplementary figure 2 and Fig. 5 – figure supplement 1). Relative values shown in Fig. 2 – supplementary figure 2 and Fig. 5 – figure supplement 1 were calculated by dividing each single number of formed CFUs at the different pH-values through the mean of formed CFUs at pH 5. Bacterial viability *in vivo* within the digestive tract of *C. floridanus* over time was analysed separately for the crop and midgut for *S. marcescens* and *Asaia* sp. (Fig. 2 and Fig. 5, respectively) and for *E. coli* (Fig. 5 – figure supplement 2). Zero-inflated generalized linear mixed models with negative binomial errors (package “glmmTMB”, Brooks et al., 2017) were used to model CFU number, with time after feeding as fixed predictor and ant colony as random effect, except for the *E. coli* model in the crop where colony was included as fixed factor as the model did not converge with colony as a random factor. Timepoints with zero bacterial growth were again excluded in the models. Relative CFU values shown in Fig. 2, Fig. 5, and Fig. 5 – supplementary figure 2 were calculated by dividing single CFU-values through the mean of CFU-values at timepoint 0h in the crop. Proportions and percentages of relative CFU change in the text are based on these relative CFU values.

## Acknowledgements

We would like to thank Robert Paxton for English grammar and style check of a pre-submission version of the manuscript, Franziska Vogel, Marvin Gilliar, and Martin Wolak for part of the data collection, Elena Crotti and Daniele Daffonchio for providing the *Asaia* strain and Martin Kaltenpoth for access to the pH microelectrode.

## Author contributions

S.T. and H.F. conceived the experiments. S.T. and M.A.M. performed the survival assays and the behavioral observations. C. H., C. T., R. B., H. F., and J. H. measured crop lumen acidity. H. F. measured pH in the midgut. M. H. and C. H. performed *in vivo* bacterial viability measurements. C. T. performed the *in vitro* bacterial growth measurements. S.T. analyzed the data and prepared the manuscript. S.T., C.H., J.H., R.B., C.T., M.H., M.A.M., R.G. and H.F. edited the manuscript.

## Competing interests

The authors declare no competing interests.

## Data and code availability

The authors declare that all data supporting the findings of this study and all code required to reproduce the analyses and figures of this study are available within the article and its supplementary information and will be made publicly available at the DRYAD digital repository upon acceptance.

---

**Fig. 1 – figure supplement 1.**
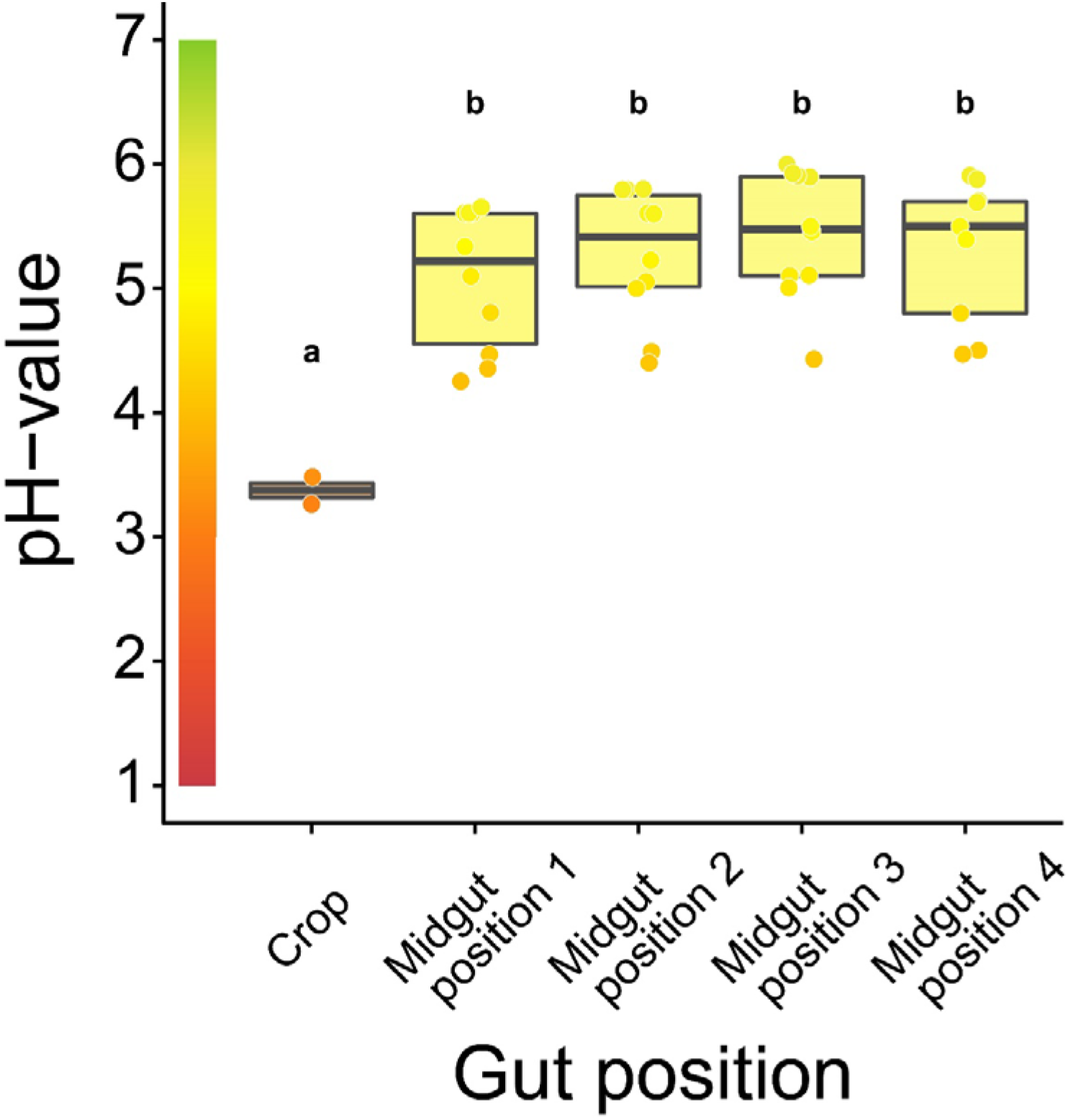
Acidity along the gastrointestinal tract of *C. floridanus*. pH-measurements 24h after access to 10% honey-water in the crop (N = 2) and directly after the proventriculus at four points along the midgut (N = 10 except position 4 with N = 9) (LMM, LR-test, χ^2^=22.152, df=4, *P* <0.001, same letters indicate *P* ≥ 0.443 and different letters indicate *P* < 0.001 in Westfall corrected post hoc comparisons). Lines and shaded boxes show median and interquartile range; points show all data. Colours in shaded rectangles near y-axis represent universal indicator pH colours. Colour filling of shaded boxes correspond to median pH colour of x-axis groups and colour filling of points correspond to universal indicator pH colours.

**Fig. 1 – figure supplement 2.**
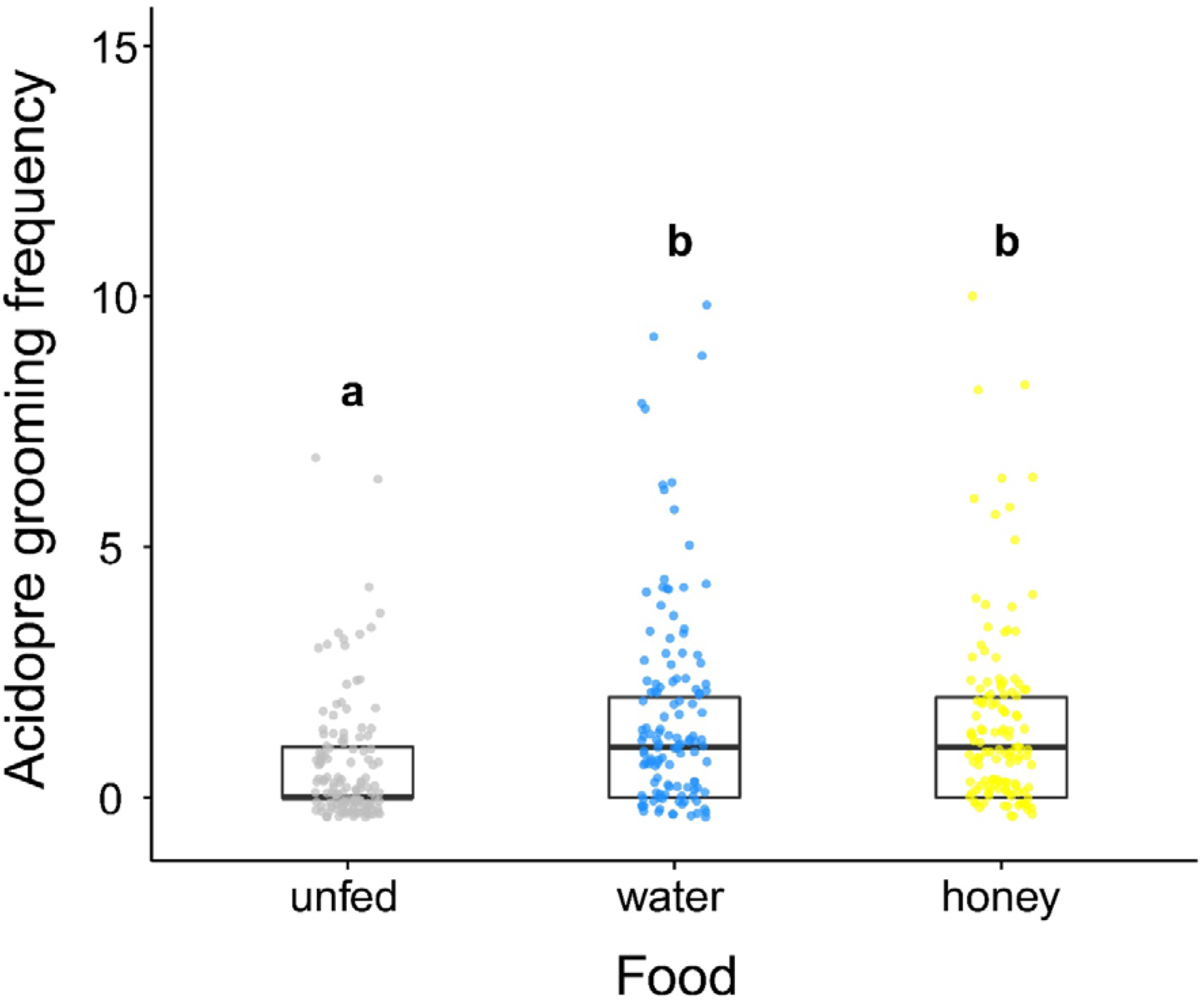
Acidopore grooming frequency of *C. floridanus* after ingestion of different food types. Frequency of acidopore grooming within 30min. after fluid ingestion (water or 10% honey water) compared to ants that did not receive any fluid (unfed) (GLMM, LR-test, χ^2^=33.526, df=2, *P* <0.001, same letters indicate *P* = 0.634 and different letters indicate *P* < 0.001 in Westfall corrected post hoc comparisons).

**Fig. 1 – figure supplement 3.**
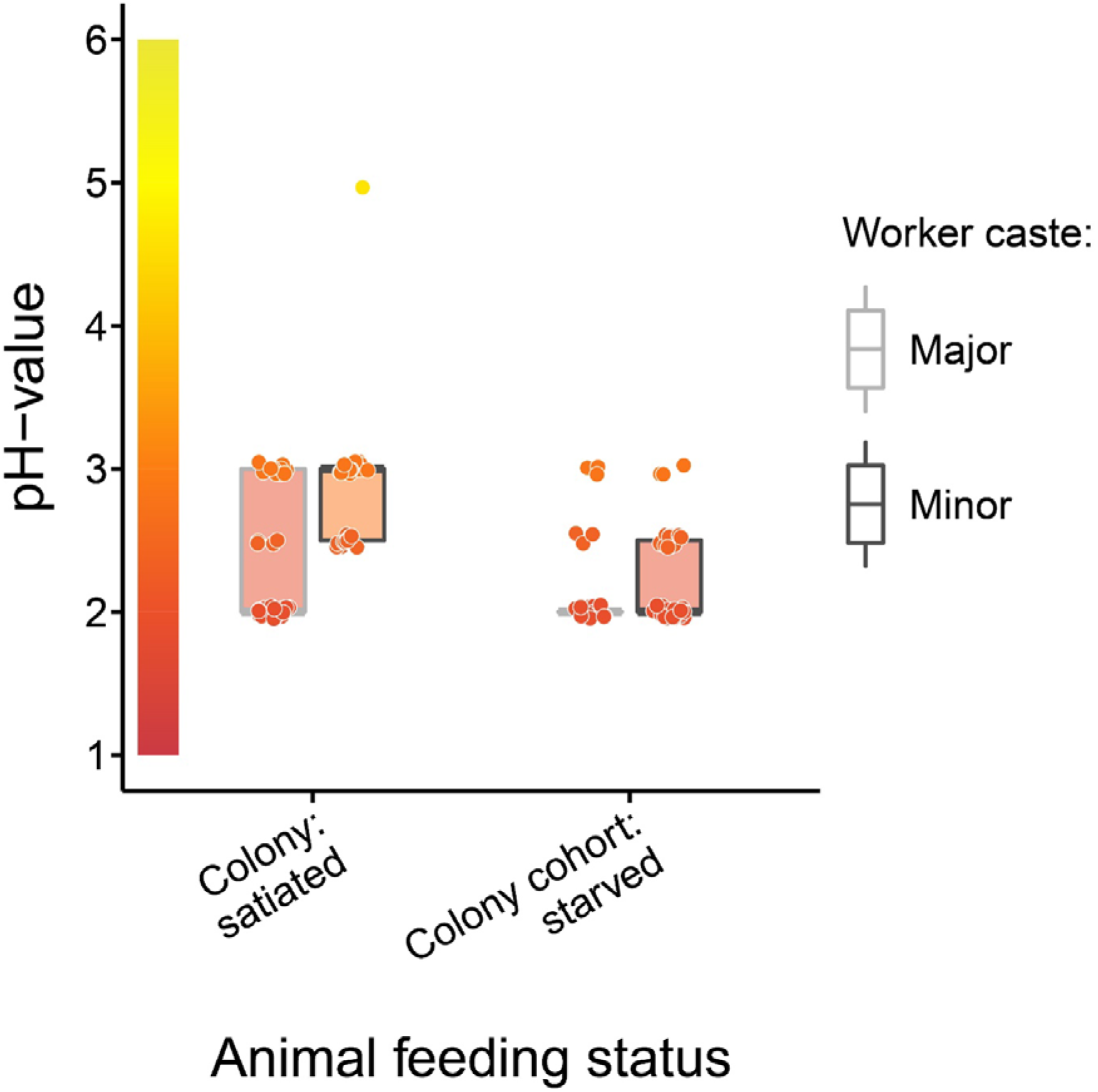
Baseline acidity of *C. floridanus* crop lumens under satiated and starved conditions. pH of crop lumens in *C. floridanus* workers (light grey: major workers, dark grey: minor workers) that were either taken directly out of a satiated colony or that were kept in cohorts of ~100 individuals under satiated conditions for three days and then starved for 24h before measuring the pH. Lines and shaded boxes show median and interquartile range; points show all data. Colours in shaded rectangles near y-axis represent universal indicator pH colours. Colour filling of shaded boxes correspond to median pH colour of x-axis groups and colour filling of points correspond to universal indicator pH colours. Border of shaded boxes represents animal caste (light grey: major workers; dark grey: minor workers).

**Fig. 2 – figure supplement 1.**
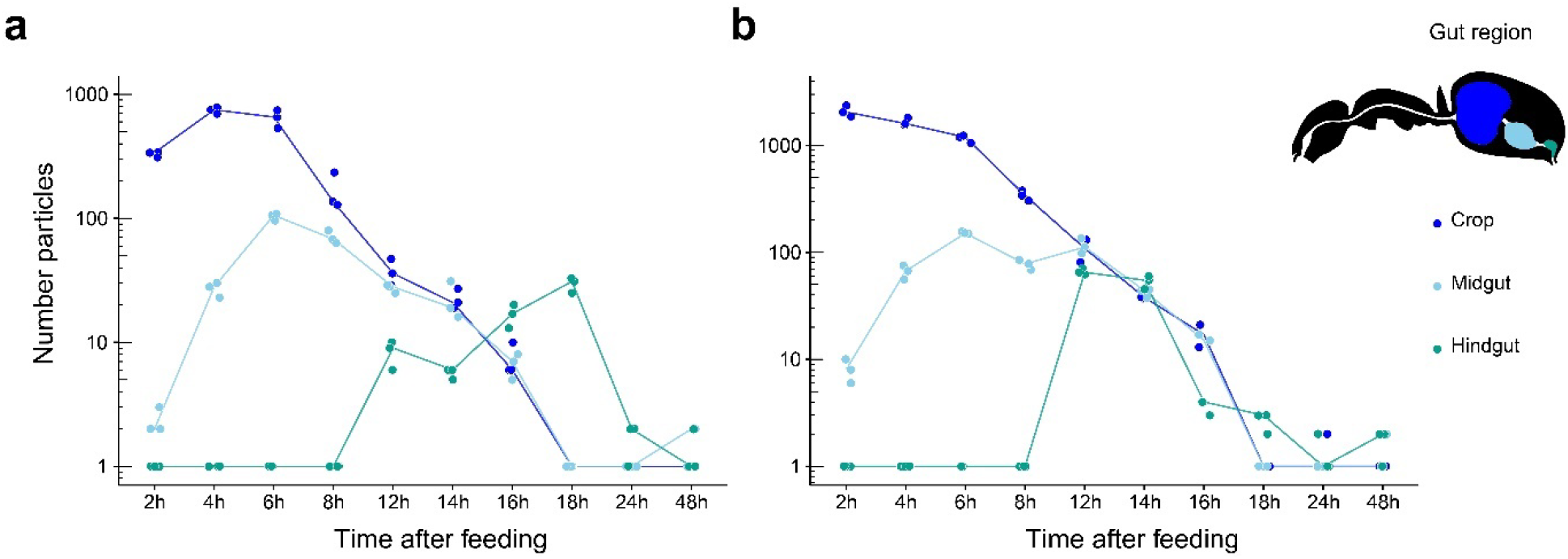
Food passage of fluorescent particles through the digestive tract of *C. floridanus*. Number of fluorescent particles on a logarithmic scale in the crop (dark blue), midgut (light blue) and hindgut (turquoise) part of the digestive tract of minor (**a**) and major (**b**) ants 2h, 4h, 6h, 8h, 12h, 14h, 16h, 18h, 24h, and 48h after feeding them a 1:1 honey-water mix with polymethylmethacrylate (PMM) particles (size ≤ 40 μm). Note that for displaying purposes and better visibility of zero values a value of one has been added to all datapoints. Points represent the number of counted particles per individual ant and lines connect the median value of particles at the different time points after feeding.

**Fig. 2 – figure supplement 2.**
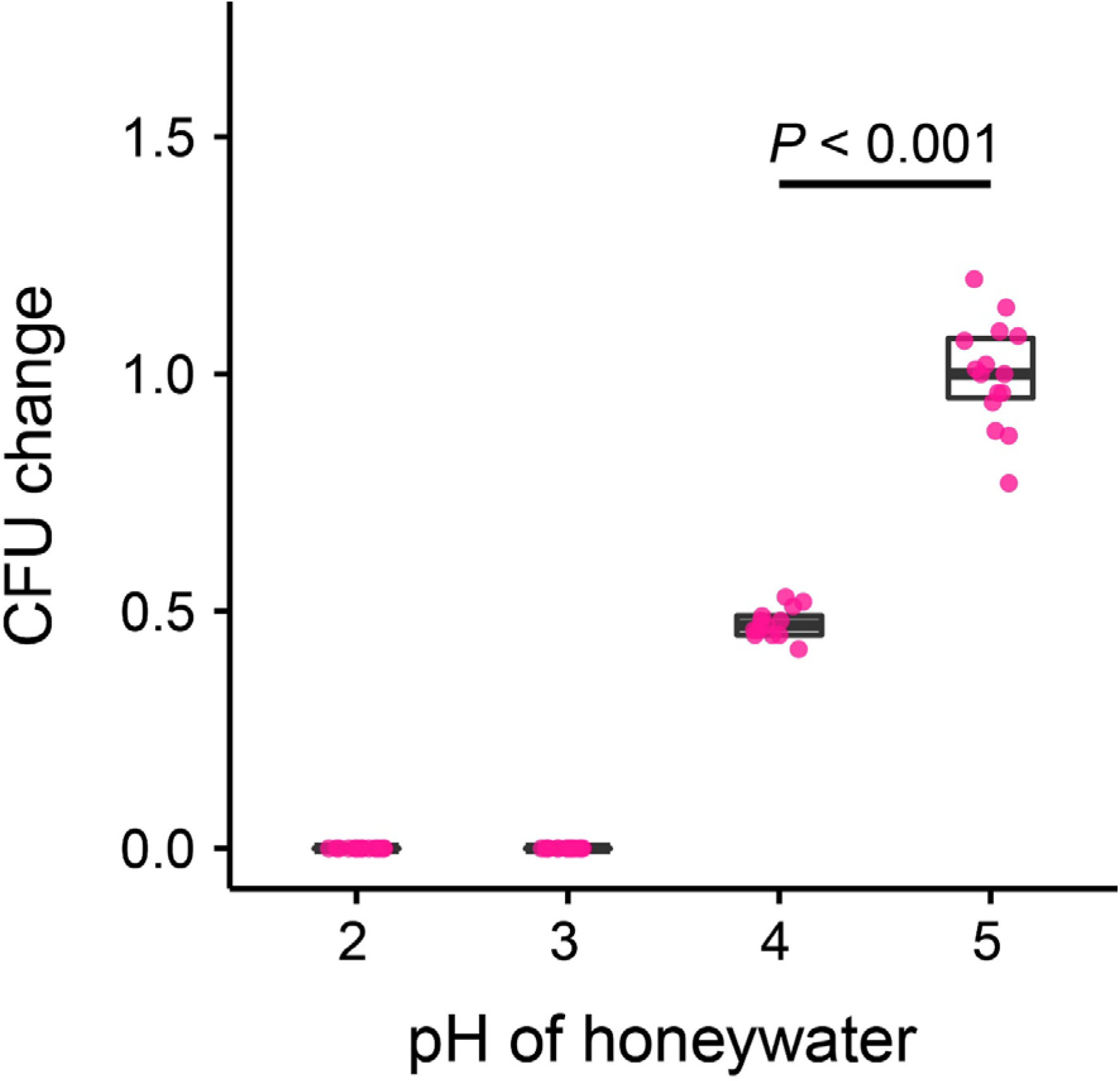
*S. marcescens* growth *in vitro*. Change in the number of CFUs relative to the mean at pH 5 (CFU change corresponds to single data CFU-value divided by the mean CFU-value at pH 5) after incubation of *Serratia marcescens* in 10% honey water (pH = 5) or in 10% honey water acidified with commercial formic acid to a pH of 4, 3 or 2 for 2h (GLM, LR-test, χ^2^ = 79.442, df = 1, *P* < 0.001). Note that pH-values with zero bacterial growth (pH 2 and 3) were excluded from the statistical model.

**Fig. 4 – figure supplement 1.**
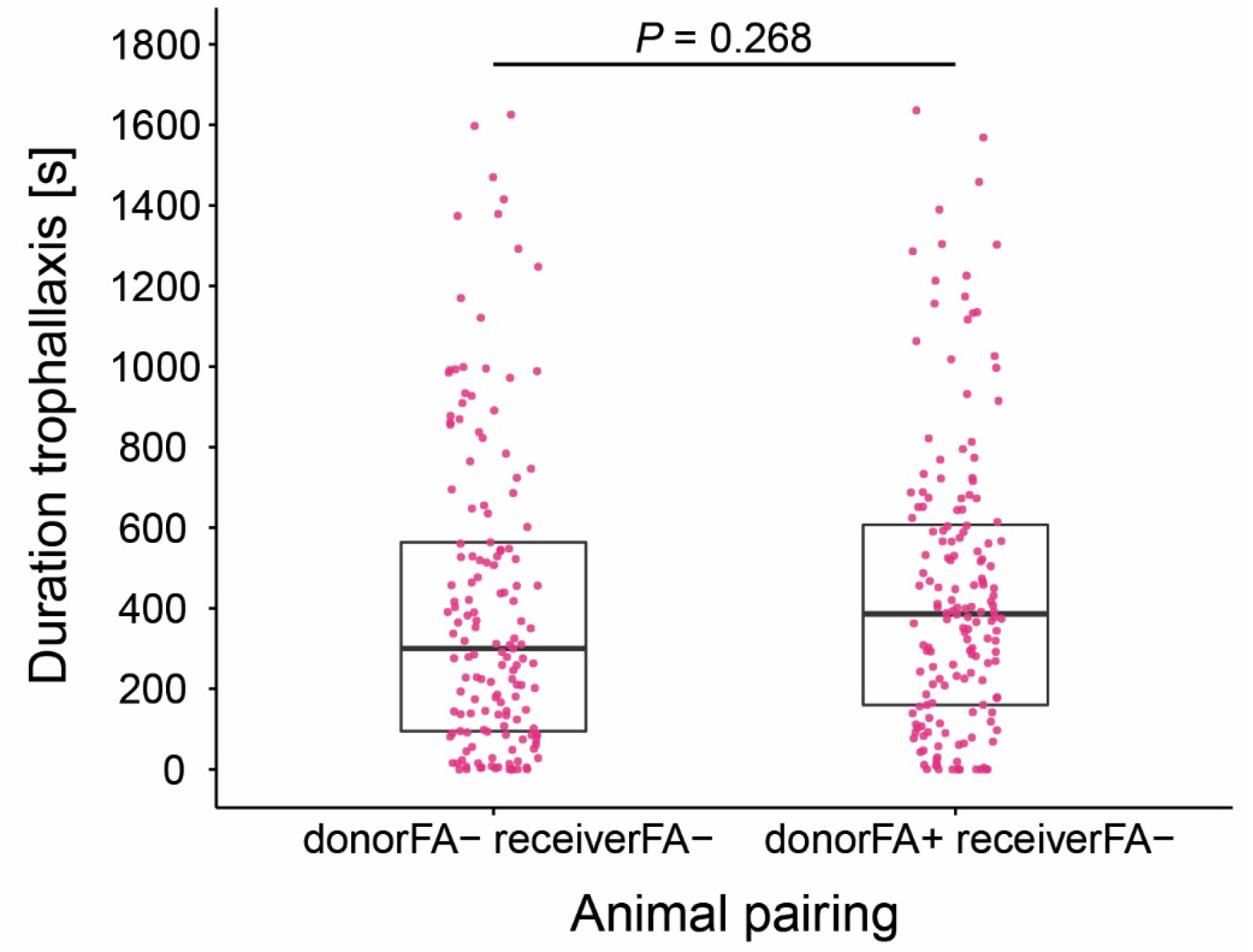
Duration of trophallaxis in donor-receiver ant pairs. Total duration of trophallaxis events within 30min. of the first bout of food exchange between donor-receiver ant-pairs (LMM, LR-test, χ^2^ = 1.23, df = 1, *P* = 0.268). Donor ants in both pairs were directly fed with *Serratia marcescens* contaminated 10% honey water and were either prevented to ingest their formic acid containing poison gland secretion (FA−) or not (FA+), while receiver ants received pathogen contaminated food only through trophallaxis with the respective donor ants and were always prevented to ingest their formic acid containing poison gland secretion (FA−).

**Fig. 5 – figure supplement 1.**
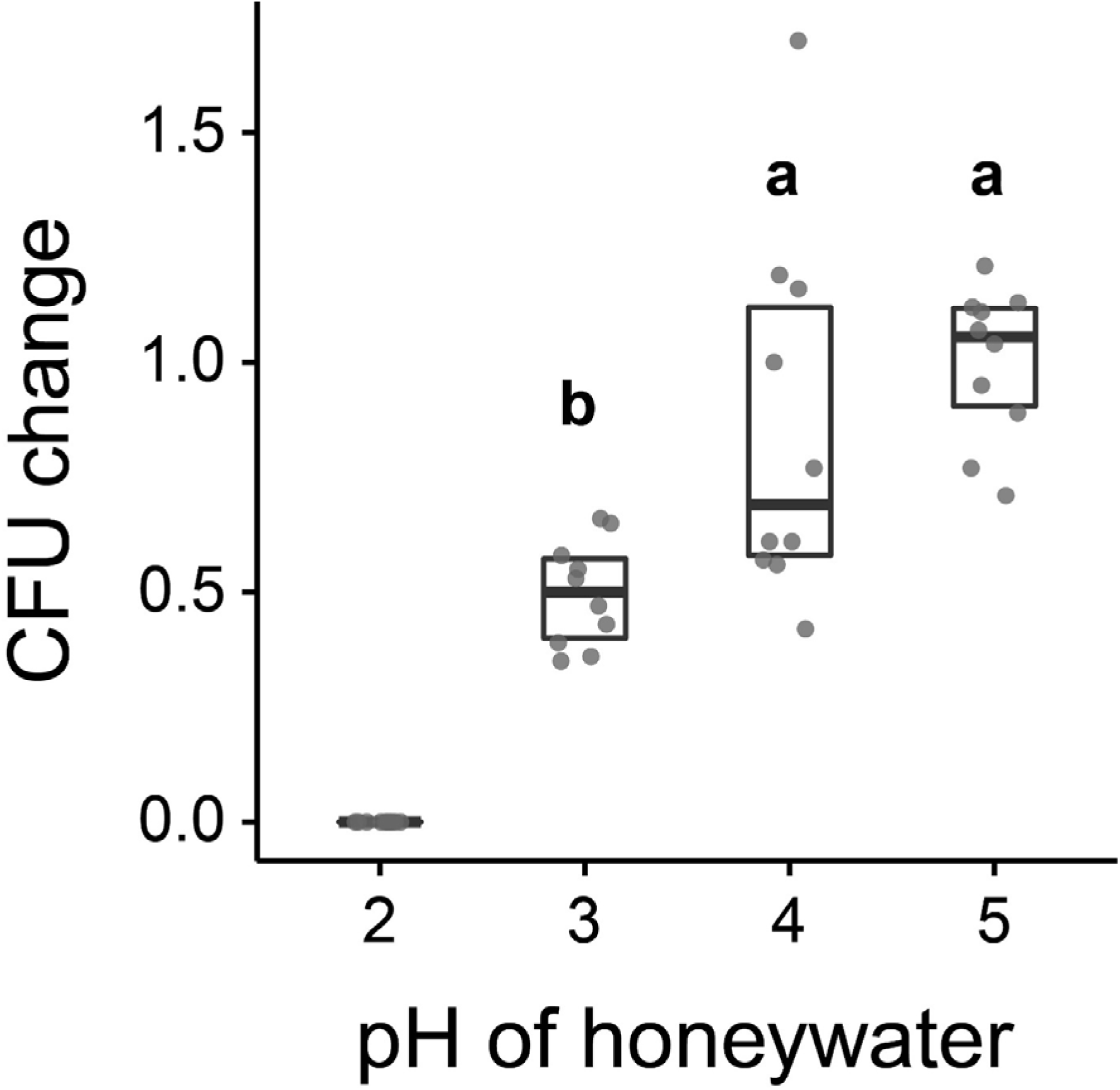
*Asaia* sp. growth *in vitro*. Change in the number of CFUs relative to the mean at pH 5 (CFU change corresponds to single data CFU-value divided by mean CFU-value at pH 5) after incubation of *Asaia* sp. in 10% honey water (pH = 5) or in 10% honey water acidified with commercial formic acid to a pH of 4, 3 or 2 for 2h (GLM, LR-test χ^2^ = 21.179, df = 2, *P* < 0.001, same letters indicate *P* = 0.234, and different letters indicate *P* < 0.001 in Westfall corrected post hoc comparisons). Note that pH-values with zero bacterial growth (pH 2) were excluded from the statistical model.

**Fig. 5 – figure supplement 2.**
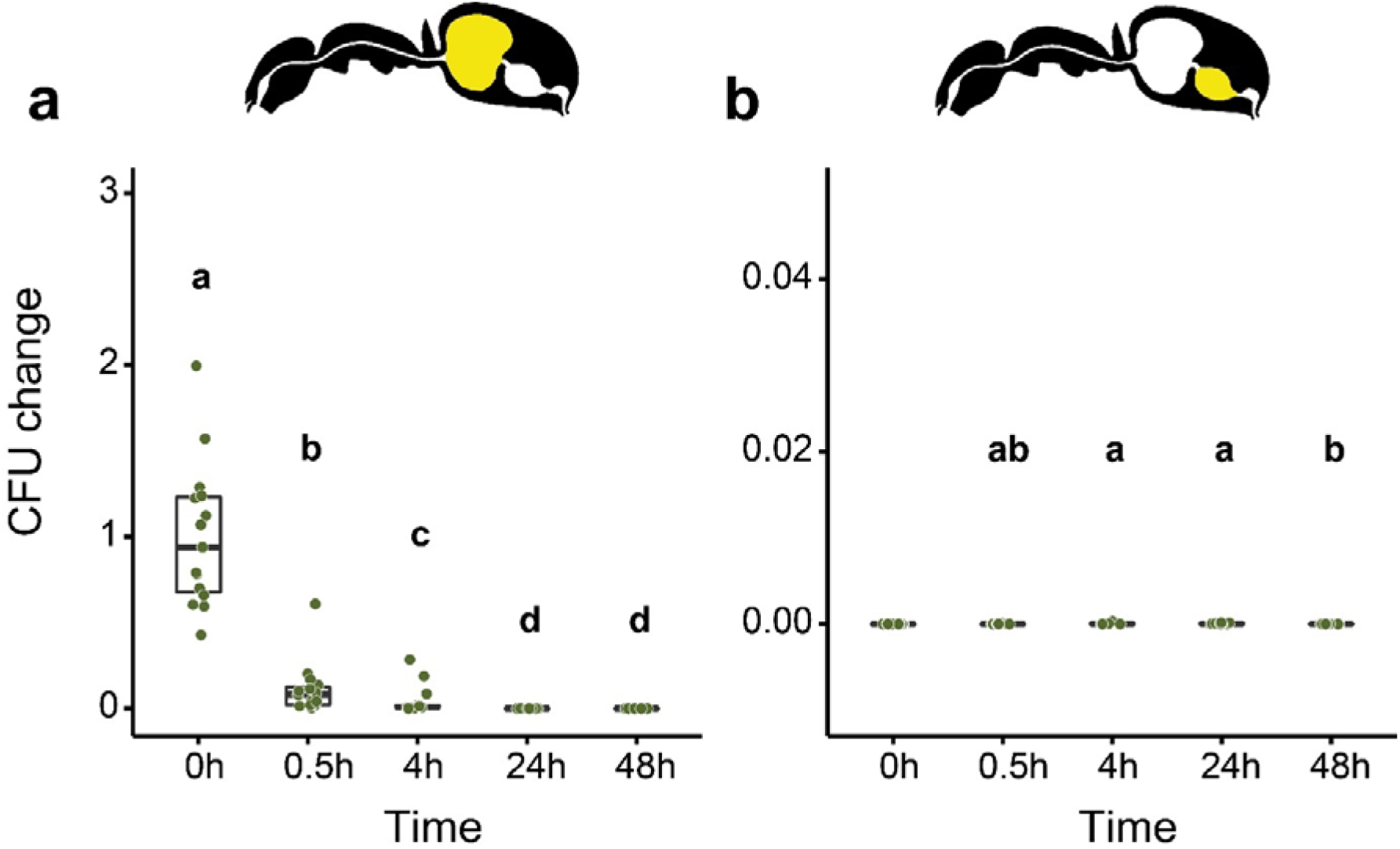
Viability of *E. coli* over time in the digestive tract of *C. floridanus* over time. Change in the number of colony forming units (CFUs) in the crop (**a**) and midgut (**b**) part of the digestive tract (yellow colour in insert) relative to mean CFU-number at 0h in the crop (CFU change corresponds to single data CFU-value divided by the mean CFU-value at 0h in the crop), 0h, 0.5h, 4h, 24h, and 48h after feeding ants 10% honey water contaminated with *Escherichia coli*. **a**, Change of *E. coli* in the crop (GLMM, LR-test, χ^2^ = 156.74, df = 4, *P* <0.001, same letters indicate *P* = 0.979 and different letters indicate *P* < 0.025 in Westfall corrected post hoc comparisons). **b**, Change of *E. coli* in the midgut (GLMM, LR-test, χ^2^ = 14.898, df = 3, *P* = 0.002, same letters indicate *P* ≥ 0.629 and different letters indicate *P* ≤ 0.038 in Westfall corrected post hoc comparisons). Note that timepoints with zero bacterial growth in the midgut (0h) were excluded from the statistical model.

## Notes

### Competing Interest Statement

The authors have declared no competing interest.

### Summary of Updates

New data have been added, the result section has been restructured and a explicit discussion section has been added

## References

Alexander, R. D. 1974. The evolution of social behavior. Annual Review of Ecology and Systematics, 5, 325–383. doi: 10.1146/annurev.es.05.110174.001545

Allaire, J. M., Crowley, S. M., Law, H. T., Chang, S. Y., Ko, H. J. & Vallance, B. A. 2018. The intestinal epithelium: central coordinator of mucosal immunity. Trends in Immunology, 39, 677–696. doi: 10.1016/j.it.2018.04.002

Anderson, K. E., Russell, J. A., Moreau, C. S., Kautz, S., Sullam, K. E., Hu, Y., Basinger, U., Mott, B. M., Buck, N. & Wheeler, D. E. 2012. Highly similar microbial communities are shared among related and trophically similar ant species. Molecular Ecology, 21, 2282–2296. doi: 10.1111/j.1365-294X.2011.05464.x

Archetti, M., Scheuring, I., Hoffman, M., Frederickson, M. E., Pierce, N. E. & Yu, D. W. 2011a. Economic game theory for mutualism and cooperation. Ecology Letters, 14, 1300–12. doi: 10.1111/j.1461-0248.2011.01697.x

Archetti, M., Ubeda, F., Fudenberg, D., Green, J., Pierce, N. E. & Yu, D. W. 2011b. Let the right one in: a microeconomic approach to partner choice in mutualisms. American Naturalist, 177, 75–85. doi: 10.1086/657622

Attygalle, A. B. & Morgan, E. D. 1984. Chemicals from the Glands of Ants. Chemical Society Reviews, 13, 245–278. doi: 10.1039/cs9841300245

Basibuyuk, H. H. & Quicke, D. L. J. 1999. Grooming behaviours in the Hymenoptera (Insecta): Potential phylogenetic significance. Zoological Journal of the Linnean Society, 125, 349–382. doi: 10.1006/zjls.1997.0167

Bates, D., Maechler, M., Bolker, B. & Walker, S. 2015. Fitting Linear mixed-effects models using lme4. Journal of Statistical Software, 67, 1–48. doi: 10.18637/jss.v067.i01

Beasley, D. E., Koltz, A. M., Lambert, J. E., Fierer, N. & Dunn, R. R. 2015. The evolution of stomach acidity and its relevance to the human microbiome. PloS One, 10, e0134116. doi: 10.1371/journal.pone.0134116

Bhatkar, A. & Whitcomb, W. H. 1971. Artificial diet for rearing various species of ants. Florida Entomologist, 53, 229–232. doi: 10.2307/3493193

Biedermann, P. H. & Kaltenpoth, M. 2014. New synthesis: the chemistry of partner choice in insect-microbe mutualisms. Journal of Chemical Ecology, 40, 99. doi: 10.1007/s10886-014-0382-8

Billen, J. 2009. Diversity and morphology of exocrine glands in ants., Ouro Preto, Brazil, Universidade Federal de Ouro Preto

Billen, J. & Sobotnik, J. 2015. Insect exocrine glands. Arthropod Structure & Development, 44, 399–400. doi: 10.1016/j.asd.2015.08.010

Billen, J. P. J. 1982. The Dufour gland closing apparatus in *Formica sanguinea* Latreille (Hymenoptera, Formicidae). Zoomorphology, 99, 235–244. doi: 10.1007/BF00312297

Blount, Z. D. 2015. The unexhausted potential of *E. coli*. eLife, 4, e05826. doi: 10.7554/eLife.05826

Blum, J. E., Fischer, C. N., Miles, J. & Handelsman, J. 2013. Frequent replenishment sustains the beneficial microbiome of *Drosophila melanogaster*. mBio, 4, e00860–13. doi: 10.1128/mBio.00860-13

Blüthgen, N., Stork, N. E. & Fiedler, K. 2004. Bottom-up control and co-occurrence in complex communities: honeydew and nectar determine a rainforest ant mosaic. Oikos, 106, 344–358. doi: 10.1111/j.0030-1299.2004.12687.x

Blüthgen, N., Verhaagh, M., Goitia, W., Jaffe, K., Morawetz, W. & Barthlott, W. 2000. How plants shape the ant community in the Amazonian rainforest canopy: the key role of extrafloral nectaries and homopteran honeydew. Oecologia, 125, 229–240. doi: 10.1007/s004420000449

Boomsma, J. J., Schmid-Hempel, P. & Hughes, W. O. H. 2005. Life histories and parasite pressure across the major groups of social insects. In: Fellowes, M. H. G. & Rolff, J. (eds.) Insect Evolutionary Ecology. Oxon, UK: CABI

Bretz, F., Hothorn, T. & Westfall, P. 2011. Multiple Comparisons using R., Boca Raton, FL, CRC Press

Brinker, P., Fontaine, M. C., Beukeboom, L. W. & Falcao Salles, J. 2019a. Host, symbionts, and the microbiome: the missing tripartite interaction. Trends in Microbiology, 27, 480–488. doi: 10.1016/j.tim.2019.02.002

Brinker, P., Weig, A., Rambold, G., Feldhaar, H. & Tragust, S. 2019b. Microbial community composition of nest-carton and adjoining soil of the ant *Lasius fuliginosus* and the role of host secretions in structuring microbial communities. Fungal Ecology, 38, 44–53. doi: 10.1016/j.funeco.2018.08.007

Broderick, N. A. & Lemaitre, B. 2012. Gut-associated microbes of *Drosophila melanogaster*. Gut Microbes, 3, 307–21. doi: 10.4161/gmic.19896

Brooks, M. E., Kristensen, K., Van Benthem, K. J., Magnusson, A., Berg, C. W., Nielsen, A., Skaug, H. J., Maechler, M. & Bolker, B. M. 2017. glmmTMB balances speed and flexibility among packages for zero-inflated generalized linear mixed modeling. The R Journal, 9, 378–400. doi:

Brown, B. P. & Wernegreen, J. J. 2016. Deep divergence and rapid evolutionary rates in gut-associated Acetobacteraceae of ants. BMC Microbiology, 16, e140. doi: 10.1186/s12866-016-0721-8

Brown, B. P. & Wernegreen, J. J. 2019. Genomic erosion and extensive horizontal gene transfer in gut-associated Acetobacteraceae. BMC Genomics, 20, e472. doi: 10.1186/s12864-019-5844-5

Brune, A. & Dietrich, C. 2015. The gut microbiota of termites: Digesting the diversity in the light of ecology and evolution. Annual Review of Microbiology, 69, 145–166. doi: 10.1146/annurev-micro-092412-155715

Brütsch, T., Jaffuel, G., Vallat, A., Turlings, T. C. & Chapuisat, M. 2017. Wood ants produce a potent antimicrobial agent by applying formic acid on tree-collected resin. Ecology and Evolution, 7, 2249–2254. doi: 10.1002/ece3.2834

Buffin, A., Denis, D., Van Simaeys, G., Goldman, S. & Deneubourg, J. L. 2009. Feeding and stocking up: Radio-labelled food reveals exchange patterns in ants. PloS One, 4, e5919. doi: 10.1371/journal.pone.0005919

Buffin, A., Mailleux, A.-C., Detrain, C. & Deneubourg, J. L. 2011. Trophallaxis in *Lasius niger*: a variable frequency and constant duration for three food types. Insectes Sociaux, 58, 177–183. doi: 10.1007/s00040-010-0133-y

Burkepile, D. E., Parker, J. D., Woodson, C. B., Mills, H. J., Kubanek, J., Sobecky, P. A. & Hay, M. E. 2006. Chemically mediated competition between microbes and animals: Microbes as consumers in food webs. Ecology, 87, 2821–2831. doi: 10.1890/0012-9658(2006)87[2821:Cmcbma]2.0.Co;2

Cannon, C. A. 1998. Nutritional ecology of the carpenter ant Camponotus pennsylvanicus (De Geer): macronutrient preference and particle consumption., Virginia, USA, Doctor Thesis of Philosophy in Entomology

Cardoza, Y. J., Klepzig, K. D. & Raffa, K. F. 2006. Bacteria in oral secretions of an endophytic insect inhibit antagonistic fungi. Ecological Entomology, 31, 636–645. doi: 10.1111/j.1365-2311.2006.00829.x

Casewell, N. R., Wuster, W., Vonk, F. J., Harrison, R. A. & Fry, B. G. 2013. Complex cocktails: The evolutionary novelty of venoms. Trends in Ecology & Evolution, 28, 219–229. doi: 10.1016/j.tree.2012.10.020

Chapman, R. F. 2013. The insects: Structure and function., Cambridge, United Kingdom, Cambridge University Press

Cherrington, C. A., Hinton, M., Mead, G. C. & Chopra, I. 1991. Organic acids: chemistry, antibacterial activity and practical applications. Advances in Microbial Physiology, 32, 87–108. doi: 10.1016/s0065-2911(08)60006-5

Chouaia, B., Gaiarsa, S., Crotti, E., Comandatore, F., Degli Esposti, M., Ricci, I., Alma, A., Favia, G., Bandi, C. & Daffonchio, D. 2014. Acetic acid bacteria genomes reveal functional traits for adaptation to life in insect guts. Genome Biology and Evolution, 6, 912–920. doi: 10.1093/gbe/evu062

Chu, H. & Mazmanian, S. K. 2013. Innate immune recognition of the microbiota promotes host-microbial symbiosis. Nature Immunology, 14, 668–75. doi: 10.1038/ni.2635

Chua, K.-O., See-Too, W.-S., Tan, J.-Y., Song, S.-L., Yong, H.-S., Yin, W.-F. & Chan, K.-G. 2020. *Oecophyllibacter saccharovorans* gen. nov. sp. nov., a bacterial symbiont of the weaver ant *Oecophylla smaragdina* with a plasmid-borne sole *rrn* operon. bioRxiv, 2020.02.15.950782. doi: 10.1101/2020.02.15.950782

Chua, K. O., Song, S. L., Yong, H. S., See-Too, W. S., Yin, W. F. & Chan, K. G. 2018. Microbial community composition reveals spatial variation and distinctive core microbiome of the weaver ant *Oecophylla smaragdina* in Malaysia. Scientific Reports, 8, e10777. doi: 10.1038/s41598-018-29159-2

Clear, M. R. & Horn, E. F. Y. 2019. The evolution of symbiotic plant-microbe signalling. Annual Plant Reviews, 2, 1–52. doi: 10.1002/9781119312994.apr0684

Cooling, M. D., Hoffmann, B. D., Gruber, M. A. M. & Lester, P. J. 2018. Indirect evidence of pathogen-associated altered oocyte production in queens of the invasive yellow crazy ant, *Anoplolepis gracilipes*, in Arnhem Land, Australia. Bulletin of Entomological Research, 108, 451–460. doi: 10.1017/S0007485317000967

Cooper, P. D. & Vulcano, R. 1997. Regulation of pH in the digestive system of the cricket, *Teleogryllus commodus* Walker. Journal of Insect Physiology, 43, 495–499. doi: 10.1016/S0022-1910(97)85495-9

Crotti, E., Damiani, C., Pajoro, M., Gonella, E., Rizzi, A., Ricci, I., Negri, I., Scuppa, P., Rossi, P., Ballarini, P., Raddadi, N., Marzorati, M., Sacchi, L., Clementi, E., Genchi, M., Mandrioli, M., Bandi, C., Favia, G., Alma, A. & Daffonchio, D. 2009. *Asaia*, a versatile acetic acid bacterial symbiont, capable of cross-colonizing insects of phylogenetically distant genera and orders. Environmental Microbiology, 11, 3252–3264. doi: 10.1111/j.1462-2920.2009.02048.x

Currie, C. R., Scott, J. A., Summerbell, R. C. & Malloch, D. 1999. Fungus-growing ants use antibiotic-producing bacteria to control garden parasites. Nature, 398, 701–704. doi: 10.1038/19519

David, L. A., Maurice, C. F., Carmody, R. N., Gootenberg, D. B., Button, J. E., Wolfe, B. E., Ling, A. V., Devlin, A. S., Varma, Y., Fischbach, M. A., Biddinger, S. B., Dutton, R. J. & Turnbaugh, P. J. 2014. Diet rapidly and reproducibly alters the human gut microbiome. Nature, 505, 559–63. doi: 10.1038/nature12820

Degnan, P. H., Lazarus, A. B., Brock, C. D. & Wernegreen, J. J. 2004. Host-symbiont stability and fast evolutionary rates in an ant-bacterium association: cospeciation of *Camponotus* species and their endosymbionts, *candidatus* Blochmannia. Systematic Biology, 53, 95–110. doi: 10.1080/10635150490264842

Demain, A. L. & Fang, A. 2000. The Natural Functions of Secondary Metabolites. In: Fiechter, A. (ed.) History of Modern Biotechnology I. Berlin, Heidelberg: Springer Berlin Heidelberg

Duarte, A., Welch, M., Swannack, C., Wagner, J. & Kilner, R. M. 2018. Strategies for managing rival bacterial communities: lessons from burying beetles. Journal of Animal Ecology, 87, 414–427. doi: 10.1111/1365-2656.12725

Eisner, T. & Happ, G. M. 1962. The infrabuccal pocket of a formicine ant: a social filtration device. Psyche, 69, 107–116. doi: 10.1155/1962/25068

Eisner, T. & Wilson, E. O. 1952. The morphology of the porventriculus of a formicine ant. Psyche, 59, 47–60. doi: 10.1155/1952/14806

Emmert, W. 1968. Die postembryonale Entwicklung sekretorischer Kopfdrüsen von *Formica pratensis* Retz. und *Apis mellifica* L. (Ins., Hym.). Zeitschrift für Morphologie der Tiere, 63, 1–62. doi: 10.1007/BF00343426

Engel, P. & Moran, N. A. 2013. The gut microbiota of insects - diversity in structure and function. FEMS Microbiology Reviews, 37, 699–735. doi: 10.1111/1574-6976.12025

Farish, D. J. 1972. The evolutionary implications of qualitative variation in the grooming behaviour of the Hymenoptera (Insecta). Animal Behaviour, 20, 662–76. doi: 10.1016/S0003-3472(72)80139-8

Favia, G., Ricci, I., Damiani, C., Raddadi, N., Crotti, E., Marzorati, M., Rizzi, A., Urso, R., Brusetti, L., Borin, S., Mora, D., Scuppa, P., Pasqualini, L., Clementi, E., Genchi, M., Corona, S., Negri, I., Grandi, G., Alma, A., Kramer, L., Esposito, F., Bandi, C., Sacchi, L. & Daffonchio, D. 2007. Bacteria of the genus *Asaia* stably associate with *Anopheles stephensi*, an Asian malarial mosquito vector. Proceedings of the National Academy of Sciences of the United States of America, 104, 9047–9051. doi: 10.1073/pnas.0610451104

Feldhaar, H., Straka, J., Krischke, M., Berthold, K., Stoll, S., Mueller, M. J. & Gross, R. 2007. Nutritional upgrading for omnivorous carpenter ants by the endosymbiont *Blochmannia*. BMC Biology, 5, 48. doi: 10.1186/1741-7007-5-48

Fernández-Marín, H., Nash, D. R., Higginbotham, S., Estrada, C., Van Zweden, J. S., D’Ettorre, P., Wcislo, W. T. & Boomsma, J. J. 2015. Functional role of phenylacetic acid from metapleural gland secretions in controlling fungal pathogens in evolutionarily derived leaf-cutting ants. Proceedings of the Royal Society of London. Series B, Biological Sciences, 282, 20150212. doi: 10.1098/rspb.2015.0212

Fernández-Marín, H., Zimmerman, J. K., Nash, D. R., Boomsma, J. J. & Wcislo, W. T. 2009. Reduced biological control and enhanced chemical pest management in the evolution of fungus farming in ants. Proceedings of the Royal Society of London. Series B, Biological sciences, 276, 2263–2269. doi: 10.1098/rspb.2009.0184

Fernández-Marín, H., Zimmerman, J. K., Rehner, S. A. & Wcislo, W. T. 2006. Active use of the metapleural glands by ants in controlling fungal infection. Proceedings of the Royal Society of London. Series B, Biological sciences, 273, 1689–95. doi: 10.1098/rspb.2006.3492

Fischbach, M. A. & Segre, J. A. 2016. Signaling in host-associated microbial communities. Cell, 164, 1288–1300. doi: 10.1016/j.cell.2016.02.037

Florez, L. V., Biedermann, P. H. W., Engl, T. & Kaltenpoth, M. 2015. Defensive symbioses of animals with prokaryotic and eukaryotic microorganisms. Natural Product Reports, 32, 904–936. doi: 10.1039/c5np00010f

Flower, N. E. & Filshie, B. K. 1976. Goblet cell membrane differentiations in the midgut of a lepidopteran larva. Journal of Cell Science, 20, 357–75. doi:

Flyg, C., Kenne, K. & Boman, H. G. 1980. Insect pathogenic properties of *Serratia marcescens*: Phage-resistant mutants with a decreased resistance to *Cecropia* immunity and a decreased virulence to *Drosophila*. Journal of General Microbiology, 120, 173–181. doi: 10.1099/00221287-120-1-173

Foster, K. R., Schluter, J., Coyte, K. Z. & Rakoff-Nahoum, S. 2017. The evolution of the host microbiome as an ecosystem on a leash. Nature, 548, 43–51. doi: 10.1038/nature23292

García-Bayona, L. & Comstock, L. E. 2018. Bacterial antagonism in host-associated microbial communities. Science, 361. doi: 10.1126/science.aat2456

Gernat, T., Rao, V. D., Middendorf, M., Dankowicz, H., Goldenfeld, N. & Robinson, G. E. 2018. Automated monitoring of behavior reveals bursty interaction patterns and rapid spreading dynamics in honeybee social networks. Proceedings of the National Academy of Sciences of the United States of America, 115, 1433–1438. doi: 10.1073/pnas.1713568115

Giannella, R. A., Zamcheck, N. & Broitman, S. A. 1972. Gastric-acid barrier to ingested microorganisms in man - studies *in-vivo* and *in-vitro*. Gut, 13, 251–256. doi: 10.1136/gut.13.4.251

Glancey, B. M., Vander Meer, R. K., Glover, A., Lofgren, C. S. & Vinson, S. B. 1981. Filtration of microparticles from liquids ingested by the red imported fire ant *Solenopsis invicta* Buren. Insectes Sociaux, 28, 295–401. doi: 10.1007/BF02224196

Gößwald, K. & Kloft, W. 1960. Untersuchungen mit radioaktiven Isotopen an Waldameisen. Entomophaga, 5, 33–41. doi: 10.1007/bf02376705

Gösswald, K. & Kloft, W. 1960a. Neuere Untersuchungen über die sozialen Wechselbeziehungen im Ameisenvolk, durchgeführt mit Radio-Isotopen. Zoologische Beiträge, 5, 519–556. doi:

Gösswald, K. & Kloft, W. 1960b. Untersuchungen mit radioaktiven Isotopen an Waldameisen. Entomophaga, 5, 33–41. doi: 10.1007/bf02376705

Grafen, A. 1990. Biological signals as handicaps. Journal of Theoretical Biology, 144, 517–46. doi: 10.1016/s0022-5193(05)80088-8

Greenberg, D. E., Porcella, S. F., Stock, F., Wong, A., Conville, P. S., Murray, P. R., Holland, S. M. & Zelazny, A. M. 2006. *Granulibacter bethesdensis* gen. nov., sp. nov., a distinctive pathogenic acetic acid bacterium in the family Acetobacteraceae. International Journal of Systematic and Evolutionary Microbiology, 56, 2609–2616. doi: 10.1099/ijs.0.64412-0

Greenwald, E., Segre, E. & Feinerman, O. 2015. Ant trophallactic networks: simultaneous measurement of interaction patterns and food dissemination. Scientific Reports, 5, 12496. doi: 10.1038/srep12496

Greenwald, E. E., Baltiansky, L. & Feinerman, O. 2018. Individual crop loads provide local control for collective food intake in ant colonies. eLife, 7, e31730. doi: 10.7554/eLife.31730

Grimont, F. & Grimont, P. A. D. 2006. The genus Serratia. In: Dworkin, M., Falkow, S., Rosenberg, E., Schleifer, K.-H. & Stackebrandt, E. (eds.) The Prokaryotes: Volume 6: Proteobacteria: Gamma Subclass. New York, NY: Springer New York

Hamilton, C., Lejeune, B. T. & Rosengaus, R. B. 2011. Trophallaxis and prophylaxis: social immunity in the carpenter ant *Camponotus pennsylvanicus*. Biology Letters, 7, 89–92. doi: 10.1098/rsbl.2010.0466

Hammer, T. J., Janzen, D. H., Hallwachs, W., Jaffe, S. P. & Fierer, N. 2017. Caterpillars lack a resident gut microbiome. Proceedings of the National Academy of Sciences of the United States of America, 114, 9641–9646. doi: 10.1073/pnas.1707186114

Harrison, J. F. 2001. Insect acid-base physiology. Annual Review of Entomology, 46, 221–250. doi: 10.1146/annurev.ento.46.1.221

Harrison, J. F., Wong, C. J. & Phillips, J. E. 1992. Recovery from acute haemolymph acidosis in unfed locusts: I. Acid transfer to the alimentary lumen is the dominant mechanism. Journal of Experimental Biology, 165, 85–96. doi:

Hartig, F. 2019. DHARMa: Residual diagnostics for hierarchical (multi-level/mixed) regression models. R package version 0.2.4. https://CRAN.R-project.org/package=DHARMa. doi:

He, H., Chen, Y. Y., Zhang, Y. L. & Wei, C. 2011. Bacteria associated with gut lumen of *Camponotus japonicus* Mayr. Environmental Entomology, 40, 1405–1409. doi: 10.1603/En11157

Hermann, H. R. & Blum, M. S. 1968. The hymenopterous poison apparatus. VI. *Camponotus pennsylvanicus* (Hymenoptera: Formicidae). Psyche, 75, Article ID 70198 doi: 10.1155/1968/70198

Hertle, R. 2005. The family of *Serratia* type pore forming toxins. Current Protein and Peptide Science, 6, 313–325. doi: 10.2174/1389203054546370

Herzner, G., Schlecht, A., Dollhofer, V., Parzefall, C., Harrar, K., Kreuzer, A., Pilsl, L. & Ruther, J. 2013. Larvae of the parasitoid wasp *Ampulex compressa* sanitize their host, the American cockroach, with a blend of antimicrobials. Proceedings of the National Academy of Sciences of the United States of America, 110, 1369–1374. doi: 10.1073/pnas.1213384110

Herzner, G. & Strohm, E. 2007. Fighting fungi with physics: food wrapping by a solitary wasp prevents water condensation. Current Biology, 17, R46–7. doi: 10.1016/j.cub.2006.11.060

Hirshfield, I. N., Terzulli, S. & O’Byrne, C. 2003. Weak organic acids: a panoply of effects on bacteria. Science Progress, 86, 245–69. doi: 10.3184/003685003783238626

Hite, J. L., Pfenning, A. C. & Cressler, C. E. 2020. Starving the enemy? Feeding behavior shapes host-parasite interactions. Trends in Ecology & Evolution, 35, 68–80. doi: 10.1016/j.tree.2019.08.004

Hölldobler, B. & Wilson, E. O. 1990. The ants, Cambridge, MA, USA, Belknap Press

Holtof, M., Lenaerts, C., Cullen, D. & Vanden Broeck, J. 2019. Extracellular nutrient digestion and absorption in the insect gut. Cell and Tissue Research, 377, 397–414. doi: 10.1007/s00441-019-03031-9

Howard, D. F. & Tschinkel, W. R. 1981. The flow of food in colonies of the fire ant, *Solenopsis invicta*: a multifactorial study. Physiological Entomology, 6, 297–306. doi: 10.1111/j.1365-3032.1981.tb00274.x

Howden, C. W. & Hunt, R. H. 1987. Relationship between gastric-secretion and infection. Gut, 28, 96–107. doi: DOI 10.1136/gut.28.1.96

Hutkins, R. W. 2019. Microbiology and technology of fermented foods. IFT Press

Imhann, F., Bonder, M. J., Vich Vila, A., Fu, J., Mujagic, Z., Vork, L., Tigchelaar, E. F., Jankipersadsing, S. A., Cenit, M. C., Harmsen, H. J., Dijkstra, G., Franke, L., Xavier, R. J., Jonkers, D., Wijmenga, C., Weersma, R. K. & Zhernakova, A. 2016. Proton pump inhibitors affect the gut microbiome. Gut, 65, 740–8. doi: 10.1136/gutjnl-2015-310376

Imirzian, N., Zhang, Y., Kurze, C., Loreto, R. G., Chen, D. Z. & Hughes, D. P. 2019. Automated tracking and analysis of ant trajectories shows variation in forager exploration. Scientific Reports, 9, 13246. doi: 10.1038/s41598-019-49655-3

Inagaki, T. & Matsuura, K. 2018. Extended mutualism between termites and gut microbes: Nutritional symbionts contribute to nest hygiene. The Science of Nature, 105, 52. doi: 10.1007/s00114-018-1580-y

Innocent, T., Holmes, N., Bassam, M. A., Schiøtt, M., Scheuring, I., Wilkinson, B., Hutchings, M. I., Boomsma, J. J. & Yu, D. W. 2018. Experimental demonstration that screening can enable the environmental recruitment of a defensive microbiome. bioRxiv. doi: 10.1101/375634

Itoh, H., Jang, S., Takeshita, K., Ohbayashi, T., Ohnishi, N., Meng, X. Y., Mitani, Y. & Kikuchi, Y. 2019. Host-symbiont specificity determined by microbe-microbe competition in an insect gut. Proceedings of the National Academy of Sciences of the United States of America, 116, 22673–22682. doi: 10.1073/pnas.1912397116

Ivens, A. B. F., Gadau, A., Kiers, E. T. & Kronauer, D. J. C. 2018. Can social partnerships influence the microbiome? Insights from ant farmers and their trophobiont mutualists. Molecular Ecology, 27, 1898–1914. doi: 10.1111/mec.14506

Janzen, D. H. 1977. Why fruits rot, seeds mold, and meat spoils. American Naturalist, 111, 691–713. doi: 10.1086/283200

Joop, G., Roth, O., Schmid-Hempel, P. & Kurtz, J. 2014. Experimental evolution of external immune defences in the red flour beetle. Journal of Evolutionary Biology, 27, 1562–1571. doi: 10.1111/jeb.12406

Kappeler, P. M., Cremer, S. & Nunn, C. L. 2015. Sociality and health: impacts of sociality on disease susceptibility and transmission in animal and human societies. Philosophical Transactions of the Royal Society of London. Series B, Biological Sciences, 370, 20140116. doi: 10.1098/Rstb.2014.0116

Kautz, S., Rubin, B. E. R. & Moreau, C. S. 2013a. Bacterial infections across the ants: Frequency and prevalence of *Wolbachia, Spiroplasma*, and *Asaia*. Psyche, e936341. doi: 10.1155/2013/936341

Kautz, S., Rubin, B. E. R., Russell, J. A. & Moreau, C. S. 2013b. Surveying the microbiome of ants: Comparing 454 pyrosequencing with traditional methods to uncover bacterial diversity. Applied and Environmental Microbiology, 79, 525–534. doi: 10.1128/Aem.03107-12

Koelz, H. R. 1992. Gastric-acid in vertebrates. Scandinavian Journal of Gastroenterology, 27, 2–6. doi: 10.3109/00365529209095998

Kohl, E., Hölldobler, B. & Bestmann, H. J. 2001. Trail and recruitment pheromones in *Camponotus socius* (Hymenoptera: Formicidae). Chemecology, 67–73. doi: 10.1007/PL00001834

Kohlmeier, P., Hollander, K. & Meunier, J. 2016. Survival after pathogen exposure in group-living insects: don’t forget the stress of social isolation! Journal of Evolutionary Biology, 29, 1867–1872. doi: 10.1111/jeb.12916

Komagata, K., Iino, T. & Yamada, Y. 2014. The family Acetobacteraceae. In: Rosenberg, E., Delong, E. F., Lory, S., Stackebrandt, E. & Thompson, F. (eds.) The Prokaryotes: Alphaproteobacteria and Betaproteobacteria. Berlin, Heidelberg: Springer Berlin Heidelberg

Koto, A., Mersch, D., Hollis, B. & Keller, L. 2015. Social isolation causes mortality by disrupting energy homeostasis in ants. Behavioral Ecology and Sociobiology, 69, 583–591. doi: 10.1007/s00265-014-1869-6

Kwong, W. K., Medina, L. A., Koch, H., Sing, K. W., Soh, E. J. Y., Ascher, J. S., Jaffe, R. & Moran, N. A. 2017. Dynamic microbiome evolution in social bees. Science Advances, 3, e1600513. doi: 10.1126/sciadv.1600513

Lanan, M. C., Rodrigues, P. A. P., Agellon, A., Jansma, P. & Wheeler, D. E. 2016. A bacterial filter protects and structures the gut microbiome of an insect. The ISME Journal, 10, 1866–1876. doi: 10.1038/ismej.2015.264

Leboeuf, A. C., Waridel, P., Brent, C. S., Goncalves, A. N., Menin, L., Ortiz, D., Riba-Grognuz, O., Koto, A., Soares, Z. G., Privman, E., Miska, E. A., Benton, R. & Keller, L. 2016. Oral transfer of chemical cues, growth proteins and hormones in social insects. eLife, 5, e20375. doi: 10.7554/eLife.20375

Little, A. E., Murakami, T., Mueller, U. G. & Currie, C. R. 2006. Defending against parasites: fungus-growing ants combine specialized behaviours and microbial symbionts to protect their fungus gardens. Biology Letters, 2, 12–16. doi: 10.1098/rsbl.2005.0371

Löfqvist, J. 1976. Formic acid and saturated hydrocarbons as alarm pheromones for the ant *Formica rufa*. Journal of Insect Physiology, 22, 1331–1346. doi: 10.1016/0022-1910(76)90155-4

Lopez, L. C., Morgan, E. D. & Brand, J. M. 1993. Hexadecanol and hexadecyl formate in the venom gland of formicine ants. Philosophical Transactions of the Royal Society B-Biological Sciences, 341, 177–180. doi: 10.1098/rstb.1993.0101

Lund, P., Tramonti, A. & De Biase, D. 2014. Coping with low pH: molecular strategies in neutralophilic bacteria. FEMS Microbiology Reviews, 38, 1091–125. doi: 10.1111/1574-6976.12076

Mamlouk, D. & Gullo, M. 2013. Acetic acid bacteria: Physiology and carbon sources oxidation. Indian Journal of Microbiology, 53, 377–384. doi: 10.1007/s12088-013-0414-z

Markin, G. P. 1970. Food distribution within laboratory colonies of the argentine ant,*Tridomyrmex humilis* (Mayr). Insectes Sociaux, 17, 127–157. doi: 10.1007/BF02223074

Martinsen, T. C., Bergh, K. & Waldum, H. L. 2005. Gastric juice: A barrier against infectious diseases. Basic & Clinical Pharmacology & Toxicology, 96, 94–102. doi: 10.1111/j.1742-7843.2005.pto960202.x

Matthews, P. G. D. 2017. Acid–base regulation in insect haemolymph. In: Weihrauch, D. & O’Donnell, M. (eds.) Acid-base balance and nitrogen excretion in invertebrates. Springer, Cham

Mazel, F., Davis, K. M., Loudon, A., Kwong, W. K., Groussin, M. & Parfrey, L. W. 2018. Is host filtering the main driver of phylosymbiosis across the tree of life? mSystems, 3, e00097–18. doi: 10.1128/mSystems.00097-18

Mcfall-Ngai, M., Hadfield, M. G., Bosch, T. C., Carey, H. V., Domazet-Loso, T., Douglas, A. E., Dubilier, N., Eberl, G., Fukami, T., Gilbert, S. F., Hentschel, U., King, N., Kjelleberg, S., Knoll, A. H., Kremer, N., Mazmanian, S. K., Metcalf, J. L., Nealson, K., Pierce, N. E., Rawls, J. F., Reid, A., Ruby, E. G., Rumpho, M., Sanders, J. G., Tautz, D. & Wernegreen, J. J. 2013. Animals in a bacterial world, a new imperative for the life sciences. Proceedings of the National Academy of Sciences of the United States of America, 110, 3229–36. doi: 10.1073/pnas.1218525110

Mcfrederick, Q. S., Cannone, J. J., Gutell, R. R., Kellner, K., Plowes, R. M. & Mueller, U. G. 2013. Specificity between lactobacilli and hymenopteran hosts is the exception rather than the rule. Applied and Environmental Microbiology, 79, 1803–1812. doi: 10.1128/AEM.03681-12

Miguel-Aliaga, I., Jasper, H. & Lemaitre, B. 2018. Anatomy and physiology of the digestive tract of *Drosophila melanogaster*. Genetics, 210, 357–396. doi: 10.1534/genetics.118.300224

Milan, N. F., Kacsoh, B. Z. & Schlenke, T. A. 2012. Alcohol consumption as self-medication against blood-borne parasites in the fruit fly. Current Biology, 22, 488–493. doi: 10.1016/j.cub.2012.01.045

Mirabito, D. & Rosengaus, R. B. 2016. A double-edged sword? The cost of proctodeal trophallaxis in termites. Insectes Sociaux, 63, 135–141. doi: 10.1007/s00040-015-0448-9

Moeller, A. H., Caro-Quintero, A., Mjungu, D., Georgiev, A. V., Lonsdorf, E. V., Muller, M. N., Pusey, A. E., Peeters, M., Hahn, B. H. & Ochman, H. 2016. Cospeciation of gut microbiota with hominids. Science, 353, 380–382. doi: 10.1126/science.aaf3951

Moran, N. A., Ochman, H. & Hammer, T. J. 2019. Evolutionary and ecological consequences of gut microbial communities. Annual Review of Ecology, Evolution, and Systematics, 50, 451–475. doi: 10.1146/annurev-ecolsys-110617-062453

Morgan, D. E. 2008. Chemical sorcery for sociality: exocrine secretions of ants (Hymenoptera: Formicidae). Myrmecological News, 11, 79–90. doi:

Moura-Alves, P., Puyskens, A., Stinn, A., Klemm, M., Guhlich-Bornhof, U., Dorhoi, A., Furkert, J., Kreuchwig, A., Protze, J., Lozza, L., Pei, G., Saikali, P., Perdomo, C., Mollenkopf, H. J., Hurwitz, R., Kirschhoefer, F., Brenner-Weiss, G., Weiner, J., 3RD, Oschkinat, H., Kolbe, M., Krause, G. & Kaufmann, S. H. E. 2019. Host monitoring of quorum sensing during *Pseudomonas aeruginosa* infection. Science, 366, eaaw1629. doi: 10.1126/science.aaw1629

Mueller, U. G., Gerardo, N. M., Aanen, D. K., Six, D. L. & Schultz, T. R. 2005. The evolution of agriculture in insects. Annual Review of Ecology Evolution and Systematics, 36, 563–595. doi: 10.1146/annurev.ecolsys.36.102003.152626

Muresan, C. I. & Buttstedt, A. 2019. pH-dependent stability of honey bee (*Apis mellifera*) major royal jelly proteins. Scientific Reports, 9, 9014. doi: 10.1038/s41598-019-45460-0

Mushegian, A. A. & Ebert, D. 2016. Rethinking “mutualism” in diverse host-symbiont communities. Bioessays, 38, 100–108. doi: 10.1002/bies.201500074

Nehme, N. T., Liegeois, S., Kele, B., Giammarinaro, P., Pradel, E., Hoffmann, J. A., Ewbank, J. J. & Ferrandon, D. 2007. A model of bacterial intestinal infections in *Drosophila melanogaster*. PLoS Pathogens, 3, e173. doi: 10.1371/journal.ppat.0030173

Ning, M., Yuan, M., Liu, M., Gao, Q., Wei, P., Gu, W., Wang, W. & Meng, Q. 2018. Characterization of cathepsin D from *Eriocheir sinensis* involved in *Spiroplasma eriocheiris* infection. Developmental & Comparative Immunology, 86, 1–8. doi: 10.1016/j.dci.2018.04.018

Ochman, H., Worobey, M., Kuo, C. H., Ndjango, J. B. N., Peeters, M., Hahn, B. H. & Hugenholtz, P. 2010. Evolutionary relationships of wild hominids recapitulated by gut microbial communities. PloS Biology, 8, e1000546. doi: 10.1371/journal.pbio.1000546

Ohbayashi, T., Takeshita, K., Kitagawa, W., Nikoh, N., Koga, R., Meng, X. Y., Tago, K., Hori, T., Hayatsu, M., Asano, K., Kamagata, Y., Lee, B. L., Fukatsu, T. & Kikuchi, Y. 2015. Insect’s intestinal organ for symbiont sorting. Proceedings of the National Academy of Sciences of the United States of America, 112, e5179–E5188. doi: 10.1073/pnas.1511454112

Onchuru, T. O., Martinez, A. J., Ingham, C. S. & Kaltenpoth, M. 2018. Transmission of mutualistic bacteria in social and gregarious insects. Current Opinion in Insect Science, 28, 50–58. doi: 10.1016/j.cois.2018.05.002

Onken, H. & Moffett, D. F. 2017. Acid–base loops in insect larvae with extremely alkaline midgut regions. In: Weihrauch, D. & O’Donnell, M. (eds.) Acid-base balance and nitrogen excretion in invertebrates. Springer, Cham

Osman, M. F. & Brander, J. 1961. Weitere Beiträge zur Kenntnis der chemischen Zusammensetzung des Giftes von Ameisen aus der Gattung *Formica*. Zeitschrift für Naturforschung Part B-Chemie Biochemie Biophysik Biologie und verwandten Gebiete, B 16, 749–753. doi: 10.1515/znb-1961-1108

Osman, M. F. H. & Kloft, W. 1961. Untersuchungen zur insektiziden Wirkung der verschiedenen Bestandteile des Giftes der kleinen roten Waldameise *Formica polyctena* Foerst. Insectes Sociaux, 8, 383–395. doi: 0.1007/BF02226557

Otti, O., Tragust, S. & Feldhaar, H. 2014. Unifying external and internal immune defences. Trends in Ecology & Evolution, 29, 625–634. doi: 10.1016/j.tree.2014.09.002

Oude Elferink, S. J., Krooneman, J., Gottschal, J. C., Spoelstra, S. F., Faber, F. & Driehuis, F. 2001. Anaerobic conversion of lactic acid to acetic acid and 1, 2-propanediol by *Lactobacillus buchneri*. Applied and Environmental Microbiology, 67, 125–32. doi: 10.1128/AEM.67.1.125-132.2001

Overend, G., Luo, Y., Henderson, L., Douglas, A. E., Davies, S. A. & Dow, J. A. 2016. Molecular mechanism and functional significance of acid generation in the *Drosophila* midgut. Scientific Reports, 6, 27242. doi: 10.1038/srep27242

Palmer-Young, E. C., Raffel, T. R. & Mcfrederick, Q. S. 2018. pH-mediated inhibition of a bumble bee parasite by an intestinal symbiont. Parasitology, 146, 380–388. doi: 10.1017/S0031182018001555

Perez-Cobas, A. E., Maiques, E., Angelova, A., Carrasco, P., Moya, A. & Latorre, A. 2015. Diet shapes the gut microbiota of the omnivorous cockroach *Blattella germanica*. FEMS Microbiology Ecology, 91, fiv022. doi: 10.1093/femsec/fiv022

Pull, C. D., Ugelvig, L. V., Wiesenhofer, F., Grasse, A. V., Tragust, S., Schmitt, T., Brown, M. J. & Cremer, S. 2018. Destructive disinfection of infected brood prevents systemic disease spread in ant colonies. eLife, 7, e32073. doi: 10.7554/eLife.32073

Quinlan, R. J. & Cherret, J. M. 1978. Studies on the role of the infrabuccal pocket of the leaf-cutting ant *Acromyrmex octospinosus* (Reich) (Hym., Formicidae). Insectes Sociaux, 25, 237–245. doi: 10.1007/BF02224744

Rakoff-Nahoum, S., Paglino, J., Eslami-Varzaneh, F., Edberg, S. & Medzhitov, R. 2004. Recognition of commensal microflora by toll-like receptors is required for intestinal homeostasis. Cell, 118, 229–41. doi: 10.1016/j.cell.2004.07.002

Ranger, C. M., Biedermann, P. H. W., Phuntumart, V., Beligala, G. U., Ghosh, S., Palmquist, D. E., Mueller, R., Barnett, J., Schultz, P. B., Reding, M. E. & Benz, J. P. 2018. Symbiont selection via alcohol benefits fungus farming by ambrosia beetles. Proceedings of the National Academy of Sciences of the United States of America, 115, 4447–4452. doi: 10.1073/pnas.1716852115

Ratzka, C., Liang, C., Dandekar, T., Gross, R. & Feldhaar, H. 2011. Immune response of the ant *Camponotus floridanus* against pathogens and its obligate mutualistic endosymbiont. Insect Biochemistry and Molecular Biology, 41, 529–36. doi: 10.1016/j.ibmb.2011.03.002

Ratzke, C., Denk, J. & Gore, J. 2018. Ecological suicide in microbes. Nature Ecology & Evolution, 2, 867–872. doi: 10.1038/s41559-018-0535-1

Ratzke, C. & Gore, J. 2018. Modifying and reacting to the environmental pH can drive bacterial interactions. PloS Biology, 16, e2004248. doi: 10.1371/journal.pbio.2004248

R CORE TEAM 2019. R: A language and environment for statistical computing. Vienna, Austria: R Foundation for Statistical Computing

Rebolleda-Gomez, M. & Wood, C. W. 2019. Unclear intentions: eavesdropping in microbial and plant systems. Frontiers in Ecology and Evolution, 7, 395. doi: 10.3389/fevo.2019.00385

Regnier, F. E. & Wilson, E. O. 1968. The alarm-defence system of the ant *Acanthomyops claviger*. Journal of Insect Physiology, 14, 955–970. doi: 10.1016/0022-1910(68)90006-1

Roh, S. W., Nam, Y. D., Chang, H. W., Kim, K. H., Kim, M. S., Ryu, J. H., Kim, S. H., Lee, W. J. & Bae, J. W. 2008. Phylogenetic characterization of two novel commensal bacteria involved with innate immune homeostasis in *Drosophila melanogaster*. Applied and Environmental Microbiology, 74, 6171–6177. doi: 10.1128/AEM.00301-08

Russell, J. A., Sanders, J. G. & Moreau, C. S. 2017. Hotspots for symbiosis: function, evolution, and specificity of ant-microbe associations from trunk to tips of the ant phylogeny (Hymenoptera: Formicidae). Myrmecological News, 24, 43–69. doi: 10.25849/myrmecol.news_024:043

Ryu, J. H., Kim, S. H., Lee, H. Y., Bai, J. Y., Nam, Y. D., Bae, J. W., Lee, D. G., Shin, S. C., Ha, E. M. & Lee, W. J. 2008. Innate immune homeostasis by the homeobox gene *caudal* and commensal-gut mutualism in *Drosophila*. Science, 319, 777–782. doi: 10.1126/science.1149357

Salem, H., Florez, L., Gerardo, N. & Kaltenpoth, M. 2015. An out-of-body experience: the extracellular dimension for the transmission of mutualistic bacteria in insects. Proceedings of the Royal Society B-Biological Sciences, 282, 20142957. doi: 10.1098/rspb.2014.2957

Scheuring, I. & Yu, D. W. 2012. How to assemble a beneficial microbiome in three easy steps. Ecology Letters, 15, 1300–1307. doi: 10.1111/j.1461-0248.2012.01853.x

Schmidt, J. O. 1986. Chemistry, pharmacology, and chemical ecology of ant venoms. In: Piek, T. (ed.) Venoms of the Hymenoptera: biochemical, pharmacological and behavioral aspects. Orlando, Florida: Academic Press

Schoeters, E. & Billen, J. 1996. The control apparatus of the venom gland in formicine ants (Hymenoptera: Formicidae). Netherlands Journal of Zoology, 46, 281–289. doi: 10.1163/156854295X00230

Scott, J. J., Oh, D. C., Yuceer, M. C., Klepzig, K. D., Clardy, J. & Currie, C. R. 2008. Bacterial protection of beetle-fungus mutualism. Science, 322, 63–63. doi: 10.1126/science.1160423

Sendova-Franks, A. B., Hayward, R. K., Wulf, B., Klimek, T., James, R., Planque, R., Britton, N. F. & Franks, N. R. 2010. Emergency networking: Famine relief in ant colonies. Animal Behaviour, 79, 473–485. doi: 10.1016/j.anbehav.2009.11.035

Shukla, S. P., Plata, C., Reichelt, M., Steiger, S., Heckel, D. G., Kaltenpoth, M., Vilcinskas, A. & Vogel, H. 2018a. Microbiome-assisted carrion preservation aids larval development in a burying beetle. Proceedings of the National Academy of Sciences of the United States of America, 115, 11274–11279. doi: 10.1073/pnas.1812808115

Shukla, S. P., Vogel, H., Heckel, D. G., Vilcinskas, A. & Kaltenpoth, M. 2018b. Burying beetles regulate the microbiome of carcasses and use it to transmit a core microbiota to their offspring. Molecular Ecology, 27, 1980–1991. doi: 10.1111/mec.14269

Slack, E., Hapfelmeier, S., Stecher, B., Velykoredko, Y., Stoel, M., Lawson, M. A., Geuking, M. B., Beutler, B., Tedder, T. F., Hardt, W. D., Bercik, P., Verdu, E. F., Mccoy, K. D. & Macpherson, A. J. 2009. Innate and adaptive immunity cooperate flexibly to maintain host-microbiota mutualism. Science, 325, 617–20. doi: 10.1126/science.1172747

Stroeymeyt, N., Grasse, A. V., Crespi, A., Mersch, D. P., Cremer, S. & Keller, L. 2018. Social network plasticity decreases disease transmission in a eusocial insect. Science, 362, 941–945. doi: 10.1126/science.aat4793

Strohm, E., Herzner, G., Ruther, J., Kaltenpoth, M. & Engl, T. 2019. Nitric oxide radicals are emitted by wasp eggs to kill mold fungi. eLife, 8, e43718. doi: 10.7554/eLife.43718

Stucki, D., Freitak, D., Bos, N. & Sundstrom, L. 2019. Stress responses upon starvation and exposure to bacteria in the ant *Formica exsecta*. PeerJ, 7, e6428. doi: 10.7717/peerj.6428

Tennant, S. M., Hartland, E. L., Phumoonna, T., Lyras, D., Rood, J. I., Robins-Browne, R. M. & Van Driel, I. R. 2008. Influence of gastric acid on susceptibility to infection with ingested bacterial pathogens. Infection and Immunity, 76, 639–645. doi: 10.1128/Iai.01138-07

Terra, W. R. & Ferreira 1994. Insect digestive enzymes: properties, compartmentalization and function. Comparative Biochemistry and Physiology Part B: Comparative Biochemistry, 109, 1–62. doi: 10.1016/0305-0491(94)90141-4

Therneau, T. 2019. coxme: Mixed Effects Cox Models. R package version 2.2-14. http://CRAN.R-project.org/package=coxme. doi:

Theron, M. M. & Rykers Lues, J. F. 2010. Organic acids and food preservation, Boca Raton, FL, USA, CRC Press

Tragust, S. 2016. External immune defence in ant societies (Hymenoptera: Formicidae): The role of antimicrobial venom and metapleural gland secretion. Myrmecological News, 23, 119–128. doi: 10.25849/myrmecol.news_023:119

Tragust, S., Brinker, P., Rossel, N. & Otti, O. 2020. Balancing life history investment decisions in founding ant queens. Frontiers in Ecology and Evolution, 8, 76. doi: 10.3389/fevo.2020.00076

Tragust, S., Mitteregger, B., Barone, V., Konrad, M., Ugelvig, L. V. & Cremer, S. 2013. Ants disinfect fungus-exposed brood by oral uptake and spread of their poison. Current Biology, 23, 76–82. doi: 10.1016/j.cub.2012.11.034

Traniello, J. F. A. 1977. Recruitment behavior, orientation, and the organization of foraging in the carpenter ant *Camponotus pennsylvanicus* de Geer (Hymenoptera: Formicidae). Behavioral Ecology and Sociobiology, 2, 61–79. doi: 10.1007/BF00299289

Trienens, M., Keller, N. P. & Rohlfs, M. 2010. Fruit, flies and filamentous fungi - experimental analysis of animal-microbe competition using *Drosophila melanogaster* and *Aspergillus* mould as a model system. Oikos, 119, 1765–1775. doi: 10.1111/j.1600-0706.2010.18088.x

Vander Meer, R. 2012. Ant interactions with soil organisms and associated semiochemicals. Journal of Chemical Ecology, 39, 728–745. doi: 10.1007/s10886-012-0140-8

Vander Wall, S. B. 1990. Food hoarding in animals, Chicago, Illinois, USA, University of Chicago Press

Villena, J., Kitazawa, H., Van Wees, S. C. M., Pieterse, C. M. J. & Takahashi, H. 2018. Receptors and signaling pathways for recognition of bacteria in livestock and crops: prospects for beneficial microbes in healthy growth strategies. Frontiers in Immunology, 9, 2223. doi: 10.3389/fimmu.2018.02223

Vogel, H., Shukla, S. P., Engl, T., Weiss, B., Fischer, R., Steiger, S., Heckel, D. G., Kaltenpoth, M. & Vilcinskas, A. 2017. The digestive and defensive basis of carcass utilization by the burying beetle and its microbiota. Nature Communications, 8, 15186. doi: 10.1038/ncomms15186

Watnick, P. I. & Jugder, B. E. 2020. Microbial control of intestinal homeostasis via enteroendocrine cell innate immune signaling. Trends in Microbiology, 28, 141–149. doi: 10.1016/j.tim.2019.09.005

Way, M. J. 1963. Mutualism between ants and honeydew-producing homoptera. Annual Review of Entomology, 8, 307–344. doi: DOI 10.1146/annurev.en.08.010163.001515

Williams, L. E. & Wernegreen, J. J. 2015. Genome evolution in an ancient bacteria-ant symbiosis: parallel gene loss among *Blochmannia* spanning the origin of the ant tribe Camponotini. PeerJ, 3, e881. doi: 10.7717/peerj.881

Wilson, E. O. & Eisner, T. 1957. Quantitative studies of liquid food transmission in ants. Insectes Sociaux, 4, 157–166. doi: 10.1007/BF02224149

Wolfe, A. J. 2005. The acetate switch. Microbiology and Molecular Biology Reviews, 69, 12–50. doi: 10.1128/MMBR.69.1.12-50.2005

Xiao, R., Wang, X., Xie, E., Ji, T., Li, X., Muhammad, A., Yin, X., Hou, Y. & Shi, Z. 2019. An IMD-like pathway mediates the intestinal immunity to modulate the homeostasis of gut microbiota in *Rhynchophorus ferrugineus* Olivier (Coleoptera: Dryophthoridae). Developmental & Comparative Immunology, 97, 20–27. doi: 10.1016/j.dci.2019.03.013

Yek, S. H. & Mueller, U. G. 2011. The metapleural gland of ants. Biological Reviews of the Cambridge Philosophical Society, 86, 774–91. doi: 10.1111/j.1469-185X.2010.00170.x

Zahavi, A. 1975. Mate selection-a selection for a handicap. Journal of Theoretical Biology, 53, 205–14. doi: 10.1016/0022-5193(75)90111-3

